# Panacea: a hyperpromiscuous antitoxin protein domain for the neutralisation of diverse toxin domains

**DOI:** 10.1101/2021.05.07.442387

**Authors:** Tatsuaki Kurata, Chayan Kumar Saha, Jessica A. Buttress, Toomas Mets, Tetiana Brodiazhenko, Kathryn J. Turnbull, Ololade F. Awoyomi, Sofia Raquel Alves Oliveira, Steffi Jimmy, Karin Ernits, Maxence Delannoy, Karina Persson, Tanel Tenson, Henrik Strahl, Vasili Hauryliuk, Gemma C. Atkinson

**Author notes:** corresponding authors: Gemma C. Atkinson, Vasili Hauryliuk. equal contribution (Tatsuaki Kurata, Chayan Kumar Saha).

## Abstract

Toxin-Antitoxin (TA) gene pairs are ubiquitous in microbial chromosomal genomes and plasmids, as well as bacteriophages. They act as regulatory switches, with the toxin limiting the growth of bacteria and archaea by compromising diverse essential cellular targets, and the antitoxin counteracting the toxic effect. To uncover previously uncharted TA diversity across microbes and bacteriophages, we analysed the conservation of genomic neighbourhoods using our computational tool FlaGs (for <u>Fla</u>nking <u>G</u>ene<u>s</u>), which allows high-throughput detection of TA-like operons. Focussing on the widespread but poorly experimentally characterised antitoxin domain DUF4065, our *in silico* analyses indicated that DUF4065-containing proteins serve as broadly distributed antitoxin components in putative TA-like operons with dozens of different toxic domains with multiple different folds. Given the versatility of DUF4065, we have renamed the domain to *Panacea* (and proteins containing the domain, PanA) after the Greek goddess of universal remedy. We have experimentally validated nine PanA-neutralised TA pairs. While the majority of validated PanA-neutralised toxins act as translation inhibitors or membrane disruptors, a putative nucleotide cyclase toxin from a *Burkholderia* prophage compromises replication and translation, as well as inducing RelA-dependent accumulation of the nucleotide alarmone (p)ppGpp. We find that Panacea-containing antitoxins form a complex with their diverse cognate toxins, characteristic of the direct neutralisation mechanisms employed by Type II TA systems. Finally, through directed evolution we have selected PanA variants that can neutralise non-cognate TA toxins, thus experimentally demonstrating the evolutionary plasticity of this hyperpromiscuous antitoxin domain.

**Significance:** Toxin-antitoxin systems are enigmatic and diverse elements of bacterial and bacteriophage genomes. We have uncovered remarkable versatility of an antitoxin protein domain, that has evolved to neutralise dozens of different toxin domains. We find that antitoxins carrying this domain – Panacea – form complexes with their cognate toxins, indicating a direct neutralisation mechanism, and that Panacea can be evolved to neutralise a non-cognate and non-homologous toxin with just two amino acid substitutions. This raises the possibility that this domain could be an adaptable universal, or semi-universal protein neutraliser with significant biotechnological and medical potential.

## Introduction

Toxin-antitoxin systems (TAs) are diverse two-gene elements that are widespread in plasmids and chromosomes of bacteria and archaea (1, 2), as well as in genomes of bacteriophages that prey on these microbes (3-6). The various protein toxins target different cellular core processes of the encoding cell to dramatically inhibit growth, while their cognate antitoxins efficiently neutralise the toxicity. Known TA toxins can act in a number of ways (1), commonly targeting translation by cutting or modifying the ribosome, translation factors, tRNAs or mRNAs. Similarly, antitoxins counteract the toxins through different mechanisms (1): through base-pairing of the antitoxin RNA with the toxin mRNA (Type I TA systems), direct protein-protein inhibition (Type II), inhibition of the toxin by the antitoxin RNA (Type III), or by indirect nullification of the toxicity (Type IV).

We have recently discovered a new class of toxin-antitoxin systems that employ <u>R</u>elA/<u>S</u>poT <u>h</u>omologue (RSH) enzymes – so-called <u>tox</u>ic <u>S</u>mall <u>A</u>larmone <u>S</u>ynthetases, toxSASs – as toxic enzymes to abrogate bacterial growth (4, 7). The toxicity of *Cellulomonas marina* toxSAS FaRel relies on the production of the nucleotide alarmone (pp)pApp, a pyrophosphorylated derivative synthesised from housekeeping adenosine nucleotides AMP, ADP and ATP (4). Accumulation of (pp)pApp results in dramatic depletion of ATP, which, in turn, leads to cessation of transcription followed by the inhibition of translation and replication (4, 8). Notably, (pp)pApp synthesis is not the only mechanism of toxicity employed by toxSAS: we have found that the majority of experimentally explored toxSASs, such as *Bacillus subtilis* la1a PhRel2, act as specific protein synthesis inhibitors that pyrophosphorylate the 3′-CCA end of tRNA to abrogate aminoacylation (7).

ToxSASs are neutralised by several different antitoxins that act via Type II and Type IV mechanisms. The antitoxin neutralising *B. subtilis* la1a PhRel2 (a tRNA-modifying toxSAS) belongs to a widespread domain family of unknown function designated by the Pfam database as DUF4065, where DUF stands for <u>d</u>omain of <u>u</u>nknown function (9). Clues about the roles of DUF4065 are limited; however, it is found in so called GepA (<u>G</u>enetic <u>e</u>lement <u>p</u>rotein <u>A</u>) proteins, previously associated with TA loci (10, 11), and is also present in the proteolysis-promoting SocA antitoxin of the replication inhibiting SocB toxin (12). We have earlier identified this domain in a putative alternative antitoxin to the RNAse MqsR, but this was not tested experimentally (10).

We asked whether given the broad distribution of DUF4065 across multiple phyla of bacteria and archaea, analysis of the genomic neighbourhood of DUF4065 can lead to the identification of novel TA systems. Using our tool FlaGs (13), we find that DUF4065 is the predicted antitoxin counterpart of at least 1,268 different putative TA system families corresponding to at least 88 distinct putative toxin-DUF4065 domain combinations, found in diverse bacteria, archaea and bacteriophages. While many of the toxins of these systems are related to classical TA toxins such as various mRNA interferases, Fic/Doc-type protein modification enzymes, and toxSASs, others have little similarity to known domains or proteins with solved structures. We have experimentally verified nine DUF4065-containing antitoxins as neutralisers of their cognate toxin partners. These novel toxins include translation inhibitors, membrane disruptors and a putative nucleotide cyclase that pleiotropically affects metabolism, compromising transcription and translation, as well as inducing RelA-dependent accumulation of the guanosine tetraphosphate alarmone nucleotide (p)ppGpp. Complex formation indicates DUF4065-containing antitoxins neutralise toxins via direct protein-protein interaction (that is, act as Type II TA systems), and we have identified substitutions that confer the ability of one antitoxin to neutralise a non-cognate toxin. Given the versatility of the antitoxin function of DUF4065, we have named the domain *Panacea* after the Greek goddess of universal remedy.

## Results

### The domain DUF4065 is found in diverse TA-like loci across bacteria, archaea and bacteriophages

As DUF4065 has previously been associated with TA systems (10-12), we asked whether it may constitute a widespread antitoxin domain paired in operons with novel toxin domains. To answer this, we used sensitive sequence searching combined with analysis of gene neighbourhoods using our tool FlaGs (13) (**Fig. S1**). Using the Hidden Markov Model (HMM) of the DUF4065 domain (9) to scan 20,209 genomes across cellular life and viruses, we identified 2,281 hits (**Dataset S1**) in prokaryotes and bacteriophages, comprising 27 phyla of bacteria, 3 phyla of archaea and 17 different bacteriophages (**Dataset S1**). Of those 2,281, 76 are present in complete genomes, allowing determination of whether they are chromosome- or plasmid-encoded according to the genome annotations. All but two of our identified DUF4065 homologues are chromosome-localised. The two exceptions annotated as plasmid-encoded (but may be minichromosomes) are archaeal, found in Haloarchaea (protein accessions WP_050049451.1 and WP_049938427.1). Most DUF4065-carrying taxa only carry a single homologue; 217 taxa have two, 45 have three, 14 have four, 12 have five and five have more than five. Of these five taxa, the taxon with the most DUF4065 homologues is the Mollicute bacterium “Strawberry lethal yellows phytoplasma (CPA)” strain NZSb11. This genome contains 25 DUF4065 homologues, of which three are predicted as being encoded in TA-like loci by our *in silico* analysis pipeline (see below).

Adapting FlaGs for analysing gene neighbourhood conservation, we find that around half of the identified DUF4065-containing proteins can be detected as being encoded in two-gene loci that are conserved across multiple species, reminiscent of TA systems (**Dataset S1**, representatives in **Fig. 1, Dataset S2** and **Fig. S2**). In total, we predicted 1,313 preliminarily TA (pTA)-like loci, using the criteria i) that there should be a maximum distance of 100 nucleotides between the two genes, ii) that this architecture is conserved in two or more species and iii) the conservation of the gene neighbourhood does not suggest longer operons than three genes (**Fig. S1**). We allowed three-gene architectures into our analysis as TAs can sometimes be found with a conserved third gene, such as MazG in the case of MazEF (14), chaperones in the case of tripartite toxin–<u>a</u>ntitoxin–<u>c</u>haperone (TAC) modules (15), or transcriptional regulators in the case of the paaR-paaA-parE system (3). By allowing three-part clusters, we have identified 25 clusters that are conserved as a third gene in a subset of genomes that encode a particular predicted TA pair (**Dataset S1**). We call these accessory proteins, annotations of which include DNA/nucleotide and protein/amino acid modification enzymes, helicases, proteases and nucleases. Each detected accessory third gene was only present in a small fraction of the genomes where the main TA pair was identified, suggesting that these third genes probably do not play a role in toxicity and neutralisation but are rather involved in an associated role such as phage defence.

**Figure 1.**
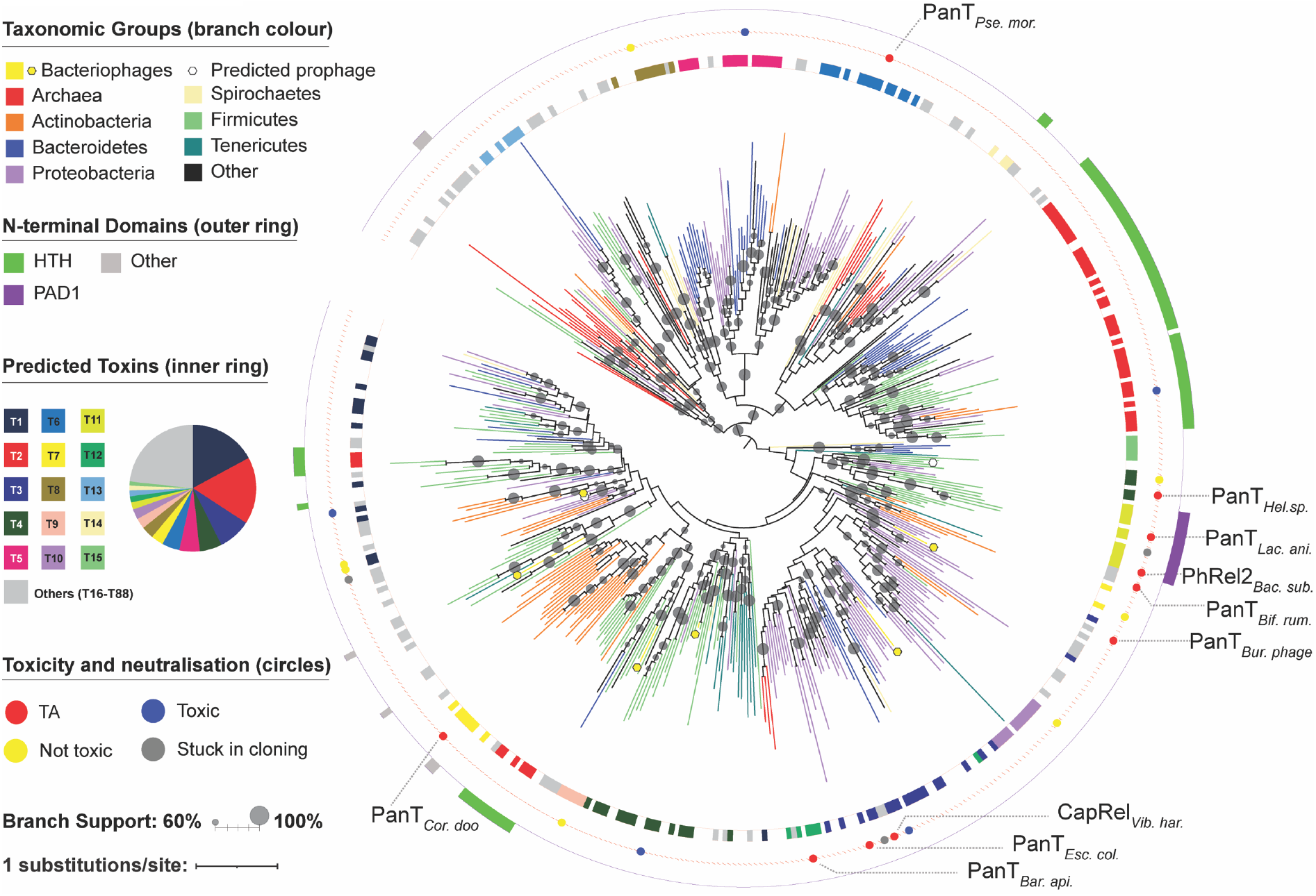
The domain DUF4065 / Panacea is found in a wide variety of TA-like loci across bacteria, archaea and bacteriophages. Branches of the IQTree maximum likelihood phylogenetic tree of representative PanA sequences are coloured by major taxonomic groupings as per the upper left key, with an additional symbol to highlight bacteriophages. Rectangles in the outer and inner rings indicate the presence and absence of N-terminal domains in the PanA sequences, and predicted associated toxin groups, respectively, according to the left-hand keys. Coloured circles between the rings indicate putative TA pairs that have been tested in toxicity neutralisation assays, and the results of those assays. Grey circles on the branches indicate branch support from IQTree ultrafast bootstrapping (42). Tree annotation was carried out with iTOL (43).

Since it is possible that some related genes are found adjacent to DUF4065-encoding genes in multiple genomes purely by chance and are not part of genuine TA systems, we set out to filter out putative “toxins” that are at risk as being spurious hits. To predict such spurious hits, we found the five closest relatives of the putative toxin in the entire set of predicted proteomes with a BlastP search, and looked for the presence of adjacent DUF4065-encoding genes (**Fig. S1**). If only the query protein is encoded adjacent to a DUF4065-encoding gene, this indicates a lack of reciprocity that suggests the potential toxin could be located in the vicinity of a DUF4065-encoding gene just by chance. From the 1,313 pTA-like loci we determined that 67 proteins (of which 39 are predicted toxins and 28 are accessory proteins) are at risk of being spurious hits (**Dataset S1**). Major classes of these spurious hits are transposases/integrases that are commonly found in TA-encoding neighbourhoods, and various ATPases that are captured into homologous clusters because of their well conserved ATP binding motifs (**Dataset S1**).

The remaining 1,268 putative TA loci that we predict to be relatively reliable correspond to 88 homologous clusters of potential toxins. We number these clusters with a T prefix; for example, SocA is in cluster T10. The vast majority of these are annotated as “hypothetical protein” as they share only weak similarity to proteins of known function. Therefore, we searched the putative toxin protein sequences against the NCBI CDD to detect the presence of known domains (**Dataset S1**). Of the 1,268 putative toxins, 938 sequences (belonging to 41 clusters) had no hit to a domain, and of the others, the most predominant domains were MqsR-like (n=90), Fic/Doc-like (n=32) and toxSAS-like (domain names NT_Pol-beta-like, RelA_SpoT and NT_Rel-Spo-like; n=31). Other known toxin domains that were represented in the CDD results were PemK (mRNAse) and ParE (DNA gyrase inhibitor). For clusters that failed to find a hit in the CDD database, HHPred (16) was run with one representative sequence, revealing additional potential homology to proteins of known structure in 28 cases (**Dataset S1**; see below for examples among our verified TAs).

The variety in the potential toxin domains suggests that the DUF4065 domain may be a universal or semi-universal antitoxin domain capable of neutralising various different toxic proteins. In light of this, we suggest renaming DUF4065 to Panacea, and abbreviate each Panacea-containing putative antitoxin and putative toxin protein as PanA, and PanT respectively. We refer to the two-gene system with the handle PanAT. In each PanAT system, the order of two genes can differ: either antitoxin first or toxin first. However, antitoxin first is the more common arrangement (943 versus 325).

Maximum Likelihood phylogenetic analysis shows the PanA tree largely does not follow taxonomic relationships, reflecting a high degree of mobility (**Fig. 1, Fig. S2 and Dataset S2**). While the deepest branches are poorly supported (not unsurprising for a small protein), there are a number of groups with medium to strong (over 60-100%) bootstrap support that include different bacterial – and sometimes archaeal – phyla. While Panacea is present broadly across prokaryotes, it does not appear to be present in eukaryotes. The only PanA we discovered in eukaryotes was in the Pharoah ant (*Monomorium pharaonic*; XP_028045404.1), and this appears to be a case of contamination as an identical sequence is found in the bacterium *Stenotrophomonas maltophilia*. Surprisingly, a strongly supported clade of PanA sequences does not necessarily mean they all share the same PanT, as shown by the inner ring in **Fig. 1** and the toxin partner swapping in focus in **Fig. 2A** and **Fig. S2**. Indeed, exchange of toxin partners within a clade appears to be frequent. We refer to this kind of domain-level partner swapping as *hyperpromiscuity*, to distinguish from the promiscuity that can be seen when one single antitoxin sequence can nullify multiple homologous toxins.

**Figure 2.**
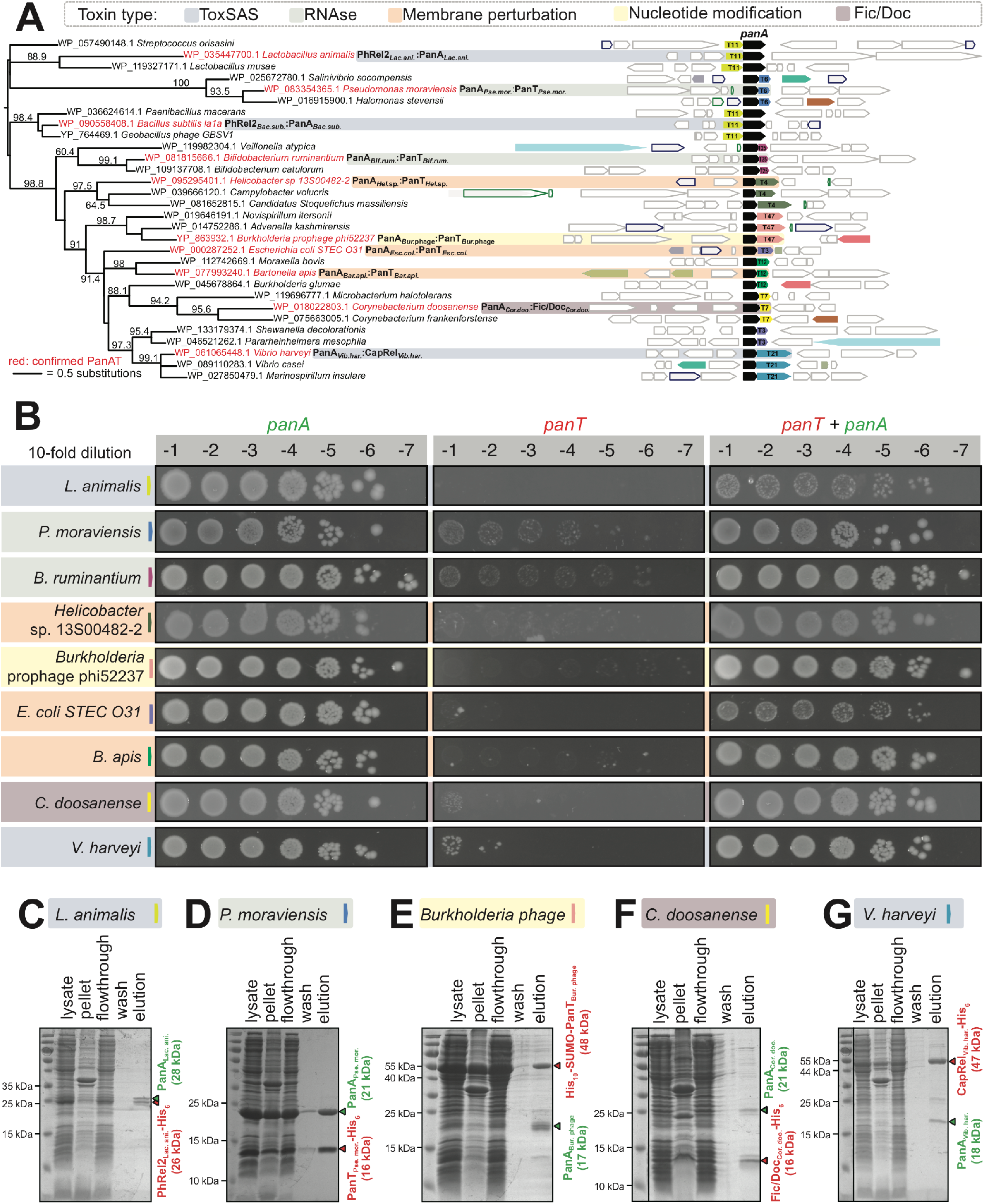
PanA antitoxins form stable complexes with evolutionarily diverse TA toxins. (**A**) The maximum likelihood tree of PanA sequences, annotated with conserved gene neighbourhoods generated with FlaGs (13). Numbers on branches show IQTree ultrafast bootstrap support (42). Genes belonging to homologous clusters are coloured the same; the PanA antitoxin is universally shown in black. Numbers on genes preceded by a T indicate toxin clusters. (**B**) Validation of *panAT* TA pairs by toxicity neutralisation assays. Overnight cultures of *E. coli* strains transformed with pBAD33 and pKK223-3 vectors or derivatives expressing putative *panT* toxins and *panA* antitoxins, correspondingly, were adjusted to OD_600_ 1.0, serially diluted from 10^1^-to 10^7^-fold and spotted on LB medium supplemented with appropriate antibiotics and inducers (0.2% arabinose for *panA* induction and 1 mM IPTG for *panT* induction). (**C**-**G**) A pull-down assay demonstrates complex formation between PanA antitoxins and PanT toxins. Untagged PanA representatives were co-expressed in *E. coli* BL21 DE3 strain together with N-terminally affinity-tagged (His_10_-SUMO in the case of *Burkholderia* prophage phi52237 PanT; His_6_ in all other cases) cognate PanT toxin. Filtered lysate was incubated with buffer-equilibrated Ni-beads, PanAT complexes were eluted with 300 mM imidazole, resolved on 15% SDS-PAGE and stained with SimplyBlue™ SafeStain.

Some – but not all – PanAs carry additional N-terminal domain regions (**Fig. 1**). Often these match a known helix-turn-helix (HTH) domain, of which a number of variations exist in the NCBI CDD. We aligned all the identified regions with hits to HTH models to make our own updated HTH model. From this, we identified HTH domains in the N-terminal regions of 343 PanA sequences (**Dataset S1**). HTH domains are often DNA-binding, are frequently found in transcription factors, and have previously been found in antitoxins, for example (17). This suggests that in some cases Panacea-domain containing antitoxins also regulate TA function at the level of transcription. Apart from HTH domains, the only widely conserved N-terminal extension appears to correspond to a new domain, which we refer to as <u>P</u>anA-<u>a</u>ssociated <u>d</u>omain 1 (PAD1) (**Fig. S3**). All but two of the TA-predicted PAD1 containing PanAs are paired with toxSAS-like toxins (the exception being putative ATPases from Clostridia (PanT group T62; **Dataset S1**). The position at the N terminus, and the presence of conserved histidines may indicate that PAD1 is a new DNA binding domain, although it has no detectable homology with any known domain. PAD1 is also present in nine Panacea-containing proteins that do not meet the criteria for TA-like loci (**Dataset S1**). In all cases where PanA contains the PAD1 domain and is in a TA-like locus, the toxin is encoded upstream of the antitoxin, the less common arrangement in the data set as a whole.

### PanA is a hyperpromiscuous antitoxin domain

Sampling broadly across PanA diversity, we selected 25 of the putative novel TAs for experimental validation in toxicity neutralisation assays (**Fig. 1** and **2A, Table 1** and **Table S1, Dataset S2**). Putative toxins and antitoxins were expressed in *Escherichia coli* strain BW25113 under the control of arabinose- and isopropyl β-D-1-thiogalactopyranoside (IPTG)-inducible promoters, respectively (18). For a gene pair to classify as a *bona fide* TA, two criteria need to be fulfilled: i) expression of the toxin should compromise *E. coli* growth and ii) co-expression the antitoxin should – either fully or partially – rescue from growth inhibition by the toxin. In addition to the PanA-neutralised PhRel2_*Bac. sub*._ toxSAS from *B. subtilis* la1a that we have validated earlier (4), we have verified here nine PanAT pairs as being genuine TA loci (**Table 1, Fig. 2B**).

**Table 1.** Summary of experimentally characterised PanAT pairs.

| Organism | Description | Toxin MOA /<br>toxin family | Toxin accession | Antitoxin<br>accession |
| --- | --- | --- | --- | --- |
| *Escherichia coli* STEC | PanA<sub>Esc. col.</sub>-PanT<sub>Esc. col.</sub> | membrane | WP_000019185.1 | WP_000287252.1 |
| *Helicobacter* sp. 13S00482-2 | PanA<sub>Hel.sp.</sub>-PanT<sub>Hel.sp.</sub> | membrane | WP_095295403.1 | WP_095295401.1 |
| *Bartonella apis* | PanA<sub>Bar. api.</sub>-PanT<sub>Bar. api.</sub> | membrane | WP_077993242.1 | WP_077993240.1 |
| *Burkholderia* prophage phi52237 | PanA<sub>Bur. phage</sub>-PanT<sub>Bur. phage</sub> | nucleotide<br>cyclase | YP_293707.1 | YP_863932.1 |
| *Bifidobacterium ruminantium* | PanA<sub>Bif. rum.</sub>-PanT<sub>Bif. rum.</sub> | RNase | WP_026646888.1 | WP_081815666.1 |
| *Pseudomonas moraviensis* | PanA<sub>Pse. mor.</sub>-PanT<sub>Pse. mor.</sub> | RNase | WP_083201923.1 | WP_083354365.1 |
| *Vibrio harveyi* | PanA<sub>Vib. har.</sub>-CapRel<sub>Vib. har.</sub> | toxSAS | WP_061065447.1 | WP_061065448.1 |
| *Bacillus subtilis* la1a | PanA<sub>Bac. sub.</sub>-PhRel2<sub>Bac. sub.</sub> | toxSAS | WP_090558406.1 | WP_090558408.1 |
| *Lactobacillus animalis* | PanA<sub>Lac. ani.</sub>-PanT<sub>Lac. ani.</sub> | toxSAS | WP_052006344.1 | WP_035447700.1 |
| *Corynebacterium doosanense* | PanA<sub>Cor. doo.</sub>-PanT<sub>Cor. doo.</sub> | Fic/Doc | WP_018022804.1 | WP_018022803.1 |

PanA-neutralised toxins from *Lactobacillus animalis* (PhRel2_*Lac. ani*._) and *Vibrio harveyi* (CapRel_*Vib. har*._) belong to two different toxSAS subfamilies, both of which we have recently shown to target translation by inhibiting tRNA aminoacylation though pyrophosphorylation of tRNA 3′-CCA end (7). Toxins from *Pseudomonas moraviensis* strain LMG 24280 (PanT_*Pse. mor*._) and *Bifidobacterium ruminantium* strain DSM 6489 (PanT_*Bif. rum*._) have no hits against the NCBI CDD, but are predicted to be structurally similar to EndoA/PemK/MazF-family RNAses with HHPred (16), and thus may act as translational inhibitors similarly to the archetypical TA toxin MazF that cleaves mRNA at ACA codons (19). The *Corynebacterium doosanense* toxin (PanT_*Cor. doo*._) is predicted to be a member of the Fic/Doc protein family, which includes the archetypal Doc TA toxin that inhibits protein synthesis by phosphorylating the essential translation elongation factor EF-Tu (20). *Burkholderia* prophage phi52237 (PanAT_*Bur*. phage_) has no detectable homology to any protein domain in the NCBI CDD. However, HHpred predicts similarity to adenylate and guanylate cyclase with 97% probability, suggesting its toxicity could be via production of a toxic cyclic nucleotide species. Finally, many of the predicted toxin genes encode putative small peptides with predicted transmembrane helices (**Fig. S4**). Of the verified TAs, the toxins with putative membrane spanning segments are those originating from *E. coli* strain STEC_O31 (PanT_*Esc. col*._), *Helicobacter* sp. 13S00482-2 (PanT_*Hel*. sp._) and *Bartonella apis* strain BBC0122 (PanT_*Bar. api*._) (**Table 1, Fig. S4**). The PanT_*Esc. col*._ and PanT_*Hel*. sp._ proteins share no detectable similarity to known toxins, but HHPred predicts weak similarity of PanT_*Bar. api*._ to the membrane-inserting toxin Fst, part of a Type I TA addiction module found on plasmids (21). The clusters containing PanT_*Esc. col*._ (T3) and PanT_*Bar. api*._ (T12) have similar sequence compositions, consisting of a charged N-terminal region, followed by a hydrophilic C terminal region where the transmembrane regions are predicted (**Fig. S2A** and **Fig. S4**). It is possible that T3 and T12 are homologous, although they are dissimilar enough that they are not clustered together by FlaGs (**Fig. S2A**). The transmembrane helices of PanT_*Hel*. sp._ are found at its C terminus, while its N-terminal region is similar to coiled coil regions found in the <u>s</u>ynaptonemal <u>c</u>omplex <u>p</u>rotein 1 (SCP-1) superfamily (22) and a Salmonella phage tail needle protein (16).

Of the potential TA pairs that were selected and could not be verified, three of the putative toxin genes could not be successfully chemically synthesised and plasmid-subcloned by the commercial provider (**Table S1**). While we can not be sure of the reason for this, it is likely that their toxicity was too severe to allow cloning in *E. coli*. Five PanTs were toxic but were not able to be rescued by their cognate PanA, and in three of these cases PanA itself was toxic (**Table S1, Figure S5A-D**). For example, while the PanA-associated mRNAse MqsR from *Herbaspirillum frisingense* GSF30 was – as we predicted earlier (10) – toxic, its toxicity was not countered by its cognate PanA when co-expressed in *E. coli* (**Figure S5C**). Finally, eight PanTs were not toxic when tested in *E. coli* – but this does not rule out the possibility of toxicity in the original host (**Table S1**).

### PanAT pairs are Type II TA systems

The Panacea domain-containing SocA antitoxin of *Caulobacter crescentus* acts as a proteolytic adaptor, bringing the toxin SocB into contact with the protease ClpPX (12). To test whether all PanAs act as such adapters, we repeated our neutralisation assays in *E. coli* strains lacking ClpPX and Lon proteases. These proteases are not necessary for neutralisation by PanA (**Fig. S6**). We therefore hypothesised that the general neutralisation mechanism of PanA is through direct binding and inhibition typical of classical Type II systems. To test this, we carried out pull-down assays using co-expressed cognate native PanA antitoxins together with N-terminally affinity-tagged with either His_6_ or His_10_-SUMO PanT toxins. We validated stable complex formation for five PanAT pairs: *L. animalis* PhRel2_*Lac. ani*._:PanA_*Lac. ani*._ (**Fig. 2C**), *P. moraviensis* PanT_*Pse. mor*._:PanA_*Pse. mor*._ (**Fig. 2D**), *Burkholderia* prophage phi52237 PanT_*Bur*. phage_: PanA_*Bur*. phage_ (**Fig. 2E**), *C. doosanense* Fic/Doc_*Cor. doo*._: PanA_*Cor. doo*._ (**Fig. 2F**) and *V. harveyi* CapRel_*Vib. har*._:PanA_*Vib. har*._ (**Fig. 2G**).

### Protein synthesis is a major target of PanT toxins

To address the molecular mechanisms of PanT toxicity, we assayed the effects of PanT expression on macromolecular synthesis by following incorporation of ^35^S methionine in proteins, ^3^H uridine in RNA and ^3^H thymidine in DNA, comparing to the effects of *E. coli* MazF RNAse as a positive control (**Fig. S7A**). As predicted, five of the identified PanT – *L. animalis* PhRel2_*Lac. ani*._ and *V. harveyi* CapRel_*Vib. har*._ toxSAS, putative RNases PanT_*Pse. mor*._ and PanT_*Bif. rum*._ and *C. doosanense* Fic/Doc toxin, Fic/Doc_*Cor. doo*._– specifically inhibit protein synthesis (**Fig. 3A-E**).

**Figure 3.**
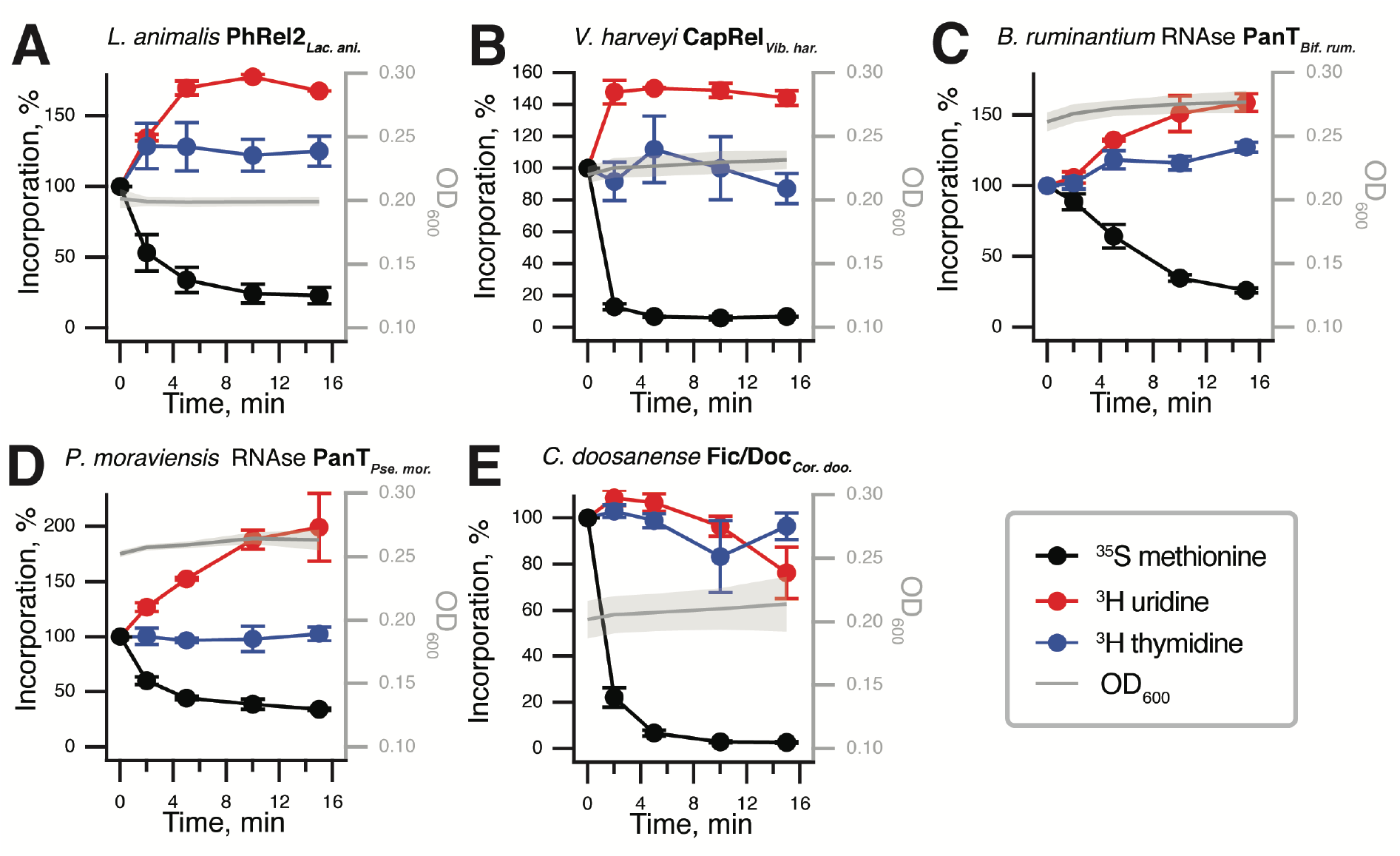
Protein synthesis is a major target of PanT toxins. Metabolic labelling assays following incorporation of ^35^S methionine (black traces), ^3^H uridine (red) and ^3^H thymidine (blue) upon expression of translation-inhibiting PanT representatives: (**A**) *L. animalis* PhRel2 and (**B**) *V. harveyi* CapRel toxSAS; putative RNases (**C**) PanT_*Bif. rum*._ and (**D**) PanT_*Pse. mor*._; (**E**) *C. doosanense* Fic/Doc toxin, Fic/Doc_*Cor. doo*._. Expression of PanTs in *E. coli* BW25113 was induced with 0.2% L-arabinose.

### *Burkholderia* prophage phi52237 PanT is a pleiotropic toxin that induces the RelA-mediated stringent response

The *Burkholderia* prophage PanT_*Bur*. phage_ toxin is unique among our verified toxins in that it predominantly inhibits transcription, with weaker effects on translation and even weaker on replication (**Fig. 4A**). The mode of inhibition is reminiscent of that of *C. marina* FaRel toxSAS (4) and *P. aeruginosa* type VI secretion system RSH effector Tas1 (7, 8) that act though production of the toxic alarmone (pp)pApp leading to dramatic depletion of ATP and GTP. Therefore, we used our HPLC-based approach to study the effects of PanT_*Bur*. phage_ toxin expression on *E. coli* nucleotide pools (23). In contrast to *C. marina* FaRel toxSAS (4), expression of PanT_*Bur*. phage_ results only in a slight decrease in GTP (**Fig. 4B**) without affecting the ATP levels (**Fig. S8A**). Surprisingly, as it is not an RSH, PanT_*Bur*. phage_ expression causes accumulation of the alarmone nucleotide ppGpp (**Fig. 4B**). This suggests that either the toxin activates cellular RelA-SpoT Homolog enzymes – given the strength of the effect, likely the stronger of the two *E. coli* (p)ppGpp synthetases, RelA – or, alternatively, the PanT_*Bur*. phage_ toxin itself is capable of producing the alarmone. No accumulation of ppGpp is detected upon PanT_*Bur*. phage_ expression in *E. coli* lacking *relA* (**Fig. 4C**); and, just as in the case of wild type, there is no effect on ATP levels in the *relA* deficient strain (**Fig. S8A**). Therefore, we conclude that the alarmone is produced by the amino acid starvation sensor RelA. To deconvolute the direct effects of *Burkholderia* prophage PanT_*Bur*. phage_ toxin on ^35^S methionine, ^3^H uridine and ^3^H thymidine incorporation from the secondary effects caused by RelA-dependent ppGpp accumulation, we performed metabolic labelling in the Δ*relA E. coli* strain (**Fig. S8C**). Just as in the wild-type strain, the main target is transcription, closely followed by translation. Thus, the growth inhibition and metabolic labelling effects observed upon PanT_*Bur*. phage_ expression are not related to ppGpp accumulation.

**Figure 4.**
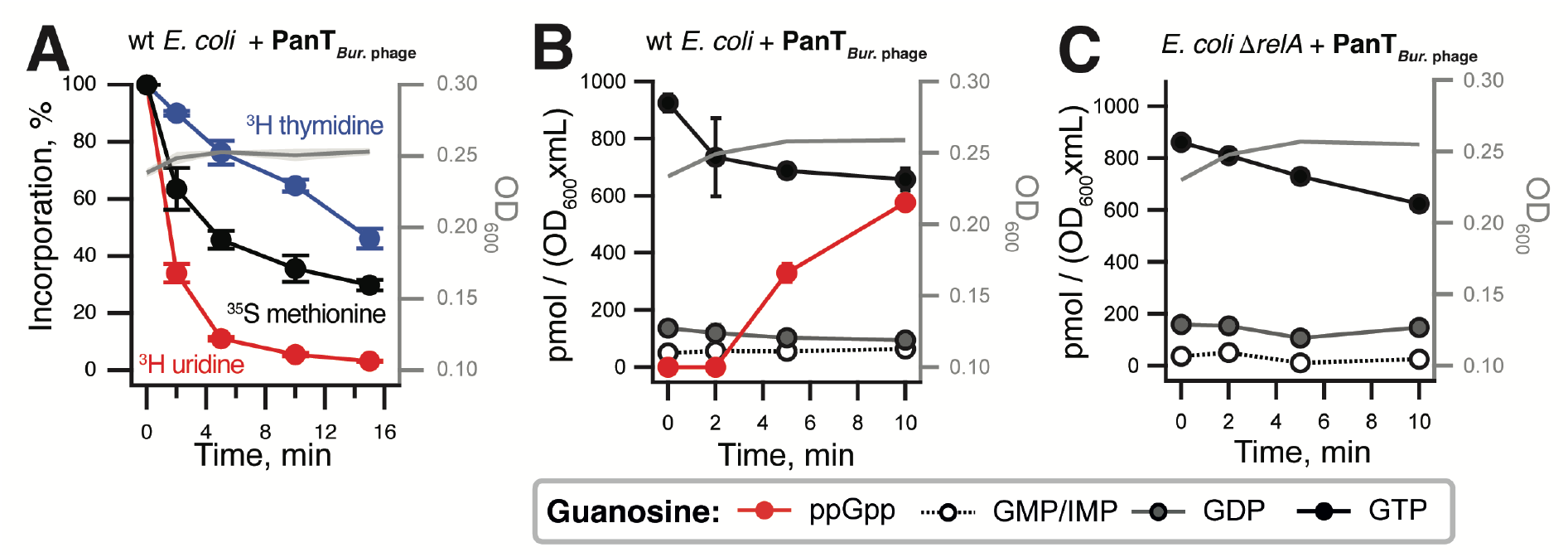
PanA-neutralised PanT_*Bur*. phage_ toxin from *Burkholderia* prophage phi52237 compromises transcription and translation, as well as inducing the RelA-mediated stringent response. (**A**) Metabolic labelling assay using wild-type *E. coli* BW25113 expressing PanT_*Bur*. phage_ toxin. (**B**,**C**) Guanosine nucleotide pools in either wild-type (**B**) or Δ*relA* (**C**) *E. coli* BW25113 expressing PanT_*Bur*. phage_ toxin. Cell cultures were grown in defined minimal MOPS medium supplemented with 0.5% glycerol at 37 °C with vigorous aeration. Expression of PanT_*Bur*. phage_ toxin was induced with 0.2% L-arabinose at the OD_600_ 0.2. Intracellular nucleotides are expressed in pmol per OD_600_ • mL as per the insert. Error bars indicate the standard error of the arithmetic mean of three biological replicates.

### The cell membrane is another major target of PanT toxins

Next, we performed ^35^S methionine, ^3^H uridine and ^3^H thymidine metabolic labelling experiments with the predicted transmembrane domain harbouring toxins PanT_*Hel*. sp._ (**Fig. 5A**), PanT_*Esc. col*._ (**Fig. 5B**) and PanT_*Bar. api*._ (**Fig. 5C**). Unlike the toxins above that predominantly target translation or transcription, expression of these toxins indiscriminately inhibited transcription, translation and DNA replication, consistent with a more general shut-down of metabolic activities caused by membrane disruption. Indeed, a comparable response was observed with induction of membrane-depolarising *E. coli* HokB TA toxin (24) (**Fig. S7B**) and treatment with the membrane-targeting inhibitor of oxidative phosphorylation <u>c</u>arbonyl <u>c</u>yanide 3-<u>c</u>hloro<u>p</u>henylhydrazone (CCCP) (**Fig. S7C**).

**Figure 5.**
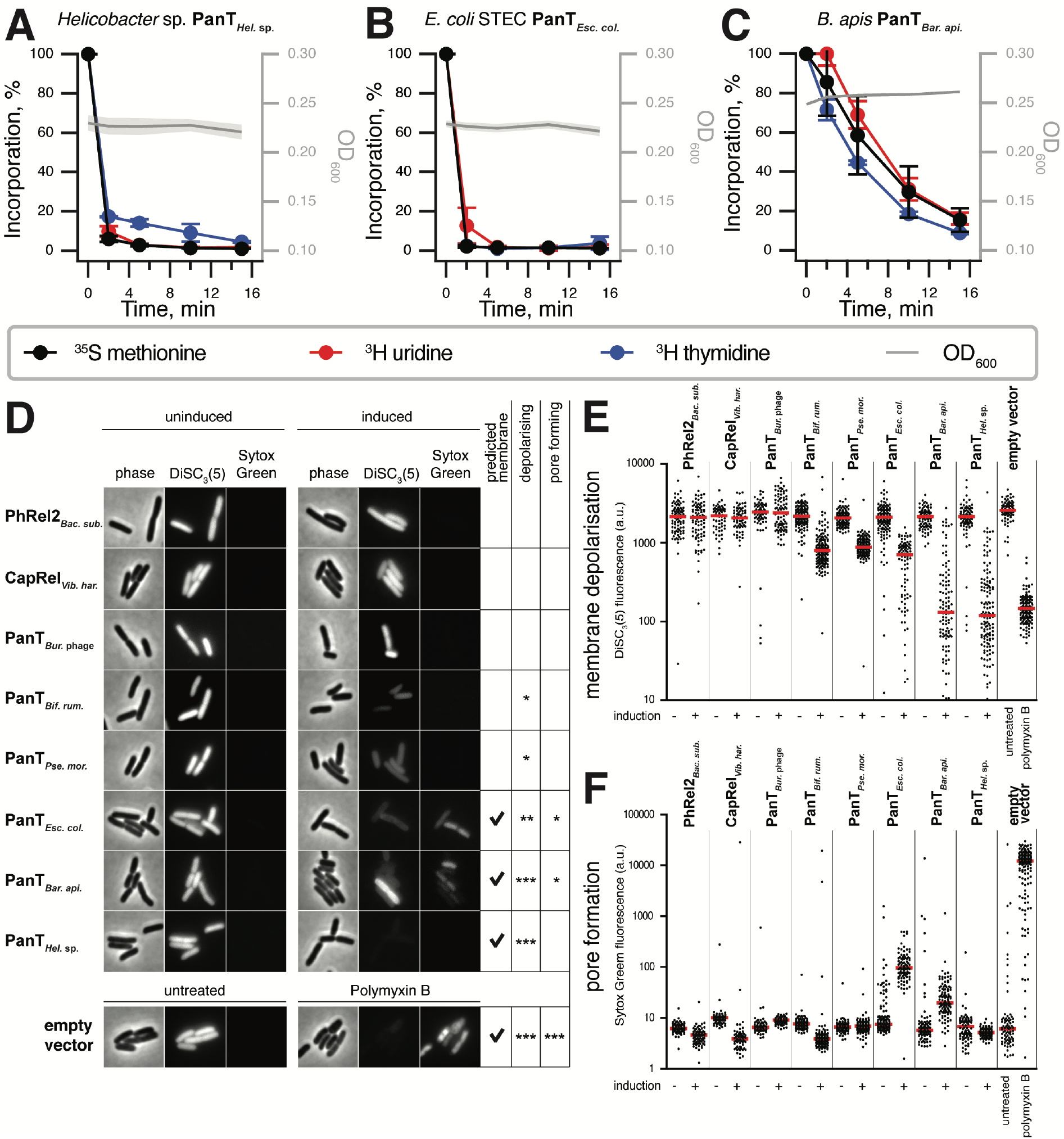
Membrane integrity is a major target of PanT toxins. (**A-C**) Metabolic labelling assays with wild-type *E. coli* BW25113 expressing (**A**) PanT_*Hel*. sp._, (**B**) PanT_*Esc. col*._ or (**C**) PanT_*Bar. api*._ toxins. (**D**) Phase-contrast (left panels) and fluorescence images (middle and right panels) of *E. coli* cells co-stained with membrane potential-sensitive dye DiSC_3_(5) and membrane permeability-indicator SYTOX Green. Depicted are representative cells carrying either an empty, or PanT-expressing vector under uninducing (no L-arabinose) or inducing (30 min induction with 0.2% L-arabinose) conditions. As a positive control, cells containing empty vector (in MOPS-glucose medium) were incubated for 15 min with membrane depolarising and pore forming antibiotic Polymyxin B. High DiSC_3_(5)-fluorescence levels indicate high membrane potential levels found in well energised, metabolically active cells. High SYTOX Green-levels, in contrast, indicate formation of pores in the inner membrane. (**E** and **F**) Quantification of (**E**) DiSC_3_(5)- and (**F**) SYTOX Green-fluorescence for individual cells from the same imaging dataset (n=92-165 cells). Median fluorescence intensity is indicated with a red line.

To directly test this hypothesis, we analysed the integrity of cell membranes upon toxin-induction using a combination of the membrane potential-sensitive dye “DiSC_3_(5)” (25) and inner membrane permeability indicator SYTOX Green (26). A strong membrane depolarisation combined with an increased SYTOX Green permeability was observed for PanT_*Bar. api*._ and PanT_*Esc. col*._ (**Fig. 5D-F**). Expression of PanT_*Hel*. sp._, in contrast, triggered strong depolarisation without an increase in SYTOX Green permeability. Thus, we conclude PanT_*Esc. col*._, PanT_*Hel*. sp._ and PanT_*Bar. api*._ exert their toxic activity through membrane depolarisation which, in the case of PanT_*Esc. col*._ and PanT_*Bar. api*._, is caused by large pore formation. Finally, weak membrane depolarisation was also observed for PanAT_*Bif. rum*._ and PanT_*Pse. mor*._ although these are not predicted to contain transmembrane helices and are instead predicted to be mRNAses. Therefore, the effect of these toxins on cell membranes is more likely to be indirect, through disturbances in respiration or central carbon metabolism. A potential membrane-spanning region is predicted for PanAT_*Bur. pro*._, although with relatively weak support (55%) (**Fig. S4D**). As this protein does not appear to affect membrane integrity, its toxicity that is particularly striking in its effect on transcription as described above, is more likely to result from its enzymatic activity, putatively cyclic nucleotide synthesis.

### While PanAs are naturally specific for their cognate PanT toxins, their PanT neutralisation spectrum can be expanded through directed evolution

We have earlier shown that Type II-antitoxins neutralising toxSAS toxins – such as *B. subtilis* la1a PanA_*Bac. sub*._ neutralising PhRel2_*Bac. sub*._ – are specific for their cognate toxins (4). PanA is clearly a versatile domain that can evolve to neutralise – and become specific for - a range of different toxin domains. Therefore, we performed an exhaustive cross-inhibition testing resulting in a 10×10 cross-neutralisation matrix (**Fig. 6A** and **Fig. S9**). A clear diagonal signal is indicative of PanA antitoxins naturally efficiently protecting only from cognate toxins – even within groups of evolutionary related toxic effectors such as toxSAS CapRel_*Vib. har*._, PhRel2_*Lac. ani*._ and PhRel2_*Bac. sub*._. Conversely, on the evolutionary timescale Panacea *does* change its toxin specificity and swaps partners, which prompts the question of whether a new specificity profile can be evolved though directed evolution.

**Figure 6.**
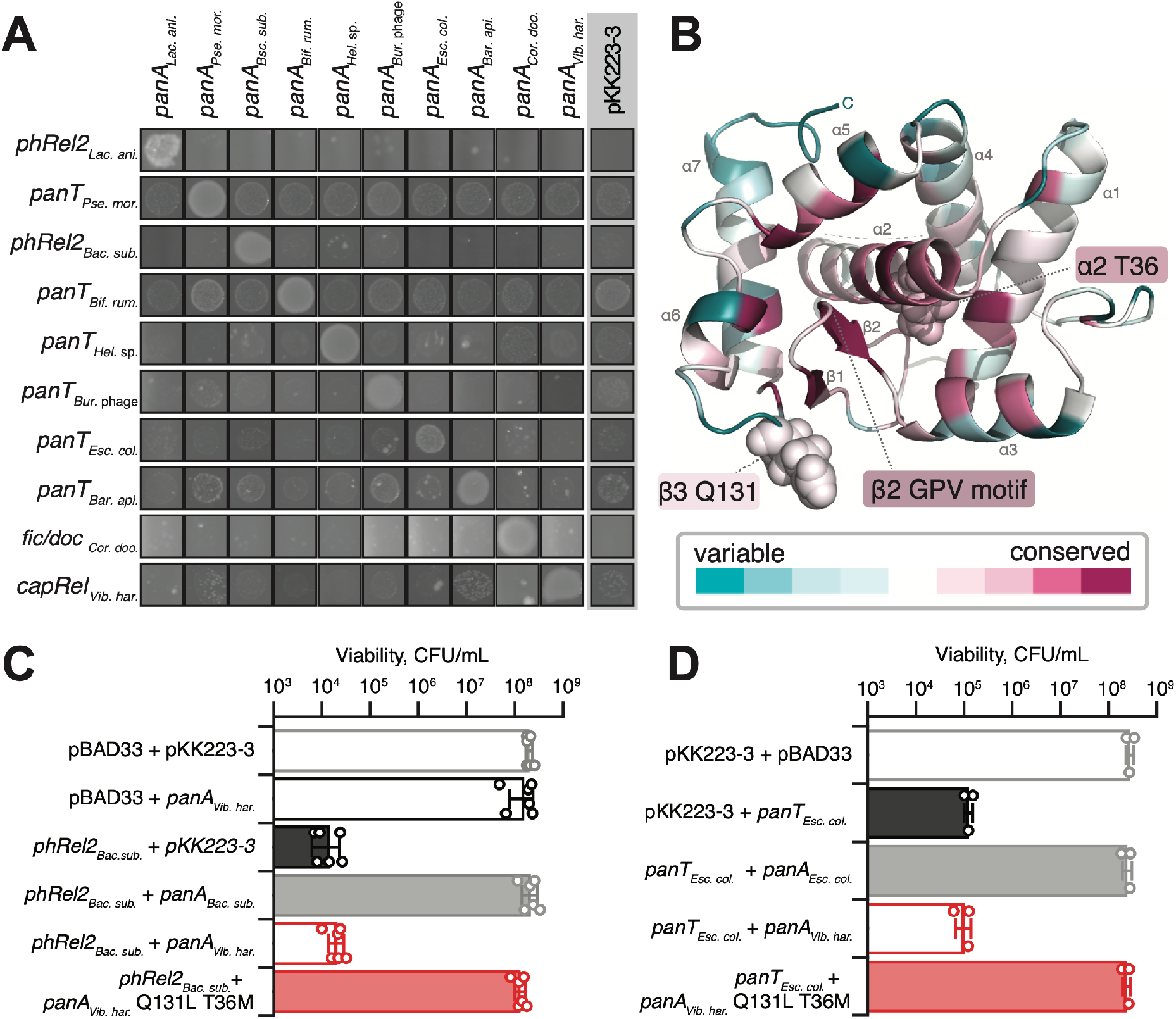
PanA specificity can be readily evolved though directed evolution. (**A**) Exhaustive cross-neutralisation testing establishes the strict specificity of PanA antitoxins towards their cognate toxins. The overnight cultures of *E. coli* strains transformed with pBAD33 and pKK223-3 vectors or derivatives thereof expressing toxin and PanA antitoxins, was adjusted to 1.0, cultures serially diluted from 10^1^- to 10^7^-fold and spotted on LB agar medium supplemented with appropriate antibiotics as well as inducers (0.2% arabinose for toxin induction and 1 mM IPTG for induction PanA variants); 10^1^-fold dilution is shown. (**B**) trRosetta-predicted structure of PanA_*Vib. har*._ antitoxin coloured by degree of conservation as per supplementary **Fig. S10**. (**C**) Neutralisation of PhRel2_*Bac. sub*._ toxin by evolved non-cognate PanA_*Vib. har*._ T36M Q131L antitoxin. To quantify the effects of PanA/PanT co-expression on bacterial viability, the overnight cultures were subdiluted, spread on the LB agar and individual colonies were counted. Analogous experiments with single-substituted T36M and Q131L PanA_*Vib. har*._ variants are shown on **Fig. S11A**. (**D**) Neutralisation of PanT_*Esc. col*._ toxin by wild-type and T36M Q131L PanA_*Vib. har*._ variants.

Structural information is useful for rationalising the effects of substitutions selected in directed evolution experiments. However, the Panacea domain is not identifiably homologous to any protein with a known structure. Therefore, we have *de novo*-predicted the structure of PanA_*Vib. har*._ using trRosetta, a deep learning-based method (27) (**Fig. 6B**). The model has a confidence categorised as “very high”, with an estimated TM-score of 0.704. The structure is comprised of a central helix (α2) surrounded by five further helices and a small three-strand beta sheet that contains a strongly conserved GPV motif in the β2 strand proximal to the central helix α2 (**Fig. 6B** and **Fig. S10**). The β3 and α2 elements are particularly well conserved in the sequence alignment (**Fig. S10)**.

Next, we targeted a pair of toxSAS:PanA TA systems with effectors belonging to two distinct toxSAS subfamilies – PhRel2 and CapRel – and screened for mutant variants of PanA _*Vib. har*._ that are able to neutralise *B. subtilis PhRel2*_*Bac. sub*._. Even though the amino acid identity between PanA _*Vib. har*._ and PanA _*Bac. sub*_ proteins is only 30-40%, just two substitutions – T36M and Q131L – were sufficient, as judged by colony counting viability testing experiments (**Fig. 6C** and **Fig. S11A**). T36 is part of the well-conserved central helix α2, while Q131 is located in a small, variable β3 strand. The β2 beta sheet containing the conserved GPV motif is sandwiched between these structural elements (**Fig. 6B** and **Fig. S10**). We asked whether individual T36M and Q131L substitutions were sufficient to elicit cross-reactivity, and concluded that they are not (**Fig. S11A**). Notably, the T36M Q131L PanA_*Vib. har*_ variant is still capable of protecting from the cognate antitoxin. However, the protection is less efficient than in the case of the cognate PanA antitoxin: the bacterial colonies are smaller, indicative of incomplete detoxification (**Fig. S11A**). Therefore, we hypothesised that the T36M Q131L double substitution does not result in specificity switching *sensu stricto*, but rather relaxes the specificity thus allowing neutralisation of non-cognate toxins. To probe this hypothesis, we tested if T36M Q131L PanA_*Vib. har*_ could protect from a noncognate cell membrane-targeting *E. coli* panT (**Fig. 6C**). We found that T36M Q131L PanA_*Vib. har*_ can, indeed, protect from PanT_*Esc. col*._ (**Fig. 6D**), although incompletely as evident from the smaller colony size (**Fig. S11B**).

## Discussion

Type II TAs are highly specific at the sequence level, however small changes can result in promiscuous intermediates allowing neutralisation of additional homologous but non-cognate toxins (28-30). Through selection experiments, we have demonstrated that via just two amino acid substitutions, Panacea-containing antitoxins can be made to neutralise not just non-cognate but *non-homologous* non-cognate toxins that have different cellular targets and mechanisms of action. This reveals a remarkable versatility of the Panacea domain. We describe the ability of an antitoxin domain to evolve to neutralise different toxin domains as *hyperpromiscuity*, distinguishing from the kind of *promiscuity* where one individual antitoxin can neutralise distinct but homologous toxins sharing the same structural fold (**Fig. 7**).

**Figure 7.**
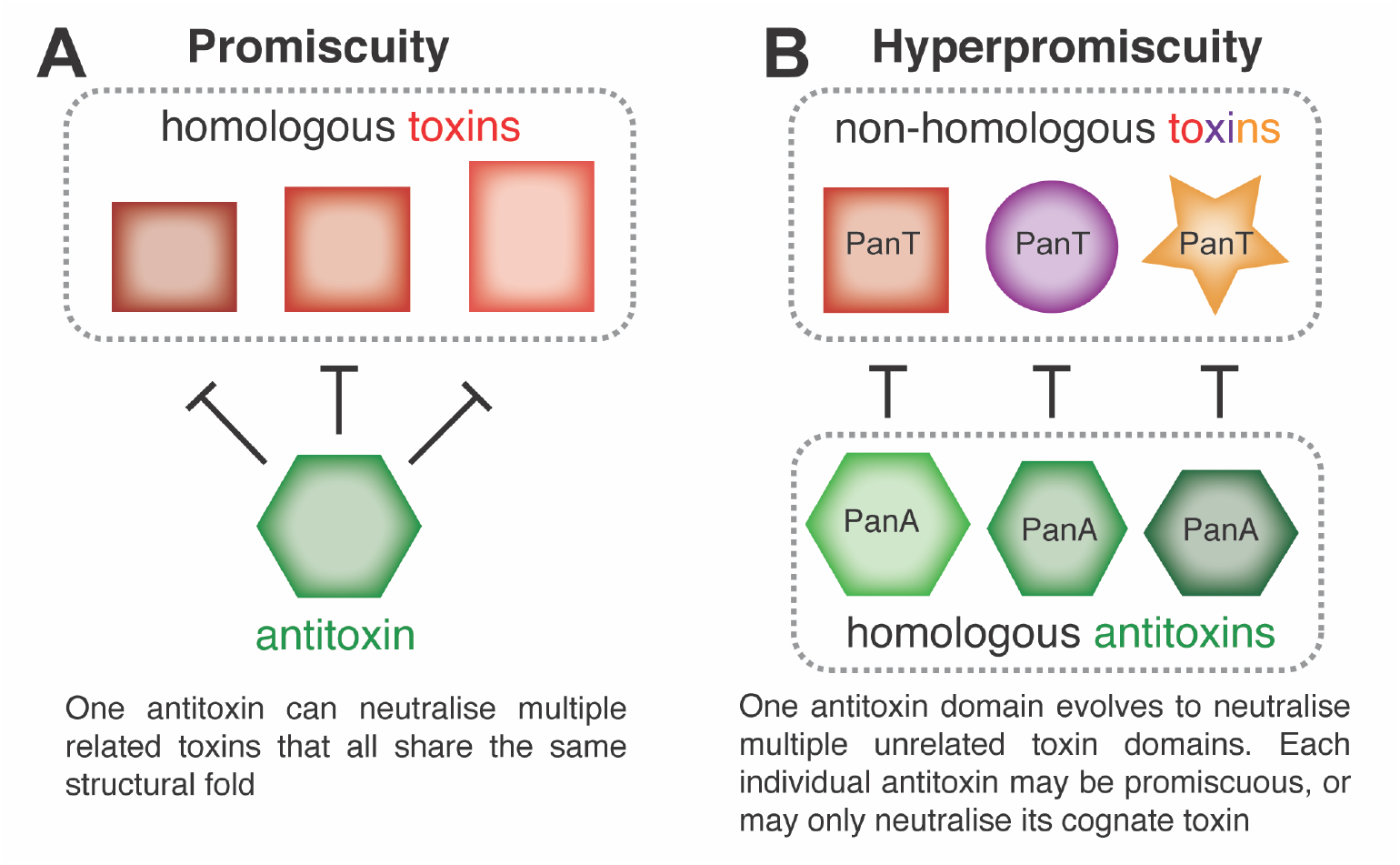
Antitoxin promiscuity *versus* hyperpromiscuity. (**A**) A promiscuous antitoxin has relaxed neutralisation specificity towards its target toxin and can neutralise a range of related toxins which all share the same structural fold. Examples include cross regulation of RelBE-like modules in *Mycobacterium tuberculosis* (44) and promiscuous ParD antitoxins generated through directed evolution that neutralise non-cognate ParE toxins (29). (**B**) A hyperpromiscuous antitoxin domain, as exemplified by Panacea, can evolve to neutralise unrelated toxins that share neither structural fold nor mechanism of action.

Other versatile antitoxin domains have also previously been observed in computational analyses to be associated with multiple toxin-like domains (11, 31, 32), indicating similar plasticity and hyperpromiscuity. One example is the PhD-related antitoxin domain found in proteins that can neutralise RelE-like mRNAses, in addition those that neutralise the EF-Tu phosphorylating toxin Doc (31). DUF4065/Panacea has previously avoided identification as a widespread antitoxin domain, despite its broad distribution. Our input set of genomes for predicting TAs is relatively small compared to the whole Genbank database. Thus, we have likely missed many additional PanAT pairs. The best way to approach this in the future is with focussed analyses of subgroups of PanA, sampling less broadly across the full diversity of Panacea, but rather focussing on more closely related PanAs, and across more closely related genomes.

A number of outstanding questions about PanA remain. Firstly, how is one single domain able to neutralise so many different toxins? The answer to this will come from structural analyses of multiple PanAs, both alone and in complex with cognate toxins. The second question is just how much of a role proteases play in the function of PanA in some species – given the previous observed function of the Panacea domain-containing antitoxin SocA in proteolytic degradation of toxin SocB in *Caulobacter* (12). Finally, the evolutionary forces that drive and enable such ready partner swapping of PanAT pairs are unclear. One answer to this is hinted at in the kinds of proteins that are encoded near PanATs (**Dataset S1**), and in the analysis of TA (though not PanAT) gene locations near recombination sites of Tn3 transposases (33). We have found that many PanATs are in close enough vicinity to transposes for them to be predicted as third component TA system genes, or even false positive potential toxins that were filtered out by our pipeline (**Dataset S1**). It is not surprising, nor a new observation that TAs can be associated with transposons; they potentially can act as addiction modules, similar to their role on plasmids (2). It is tempting to speculate that the presence of TAs near hotspots of genomic rearrangements involving transposons and prophages could lead to disruption and recombining of TA pairs.

## Materials and methods

### Identification of PanA in proteomes across the tree of life

From the NCBI genomes FTP site (ftp.ncbi.nlm.nih.gov/genomes), we downloaded 20,209 predicted proteomes, selecting all viruses, and one representative proteome per species for archaea, bacteria and eukaryotes. The full taxonomy was also retrieved from NCBI. To detect the presence of PanA across the tree of life we used the Hidden Markov Model (HMM) of the DUF4065 domain from Pfam database (9). We used HMMer v3.1b2 (34) to scan our database of proteomes with the DUF4065 HMM using thresholds set to the HMM profile’s gathering cutoffs. We found that the DUF4065 domain was present in 2,281 identified sequences. We stored the sequences, taxonomy of the source organism, domain composition in a MySQL database. We used this dataset and subsets of it for further phylogenetic analysis (see Supplementary Methods: *Representative sequence dataset assembly* and *Phylogenetic analysis*).

### Prediction of sequence features and structure

Structural modelling was carried out with the trRosetta server (27). This prediction is based on *de novo* folding, guided by deep learning restraints. Confidence in the resulting model was classified by trRosetta as “very high (with estimated TM-score=0.704).” The model was coloured by conservation using the Consurf server and an alignment of the sequences shown in **Fig. 2** (35). Transmembrane regions were predicted with the TMHMM 2.0 sever (default settings). See Supplementary Methods: *Prediction of sequence features and structure* for details of sequence analyses for prediction of protein domains, and identification of prophage-like genomic regions.

### Prediction of TA loci

Our Python tool FlaGs (13), which takes advantage of the sensitive sequence search method Jackhmmer (34), was adapted to identify conserved two- or three-gene conserved architectures that are typical of TA loci. Full details of the method are described in Supplementary Methods: *Prediction of TA loci*, with a schematic of the workflow shown in **Fig. S1**. All scripts and datasets are available at https://github.com/GCA-VH-lab/Panacea.

### Metabolic labelling with ^35^S methionine, ^3^H uridine or ^3^H thymidine

Metabolic labelling assays were performed as described previously (18). For details see Supplementary Methods: *Metabolic labelling with* ^*35*^*S methionine*, ^*3*^*H uridine or* ^*3*^*H thymidine*

### Construction of plasmids

All bacterial strains and plasmids used in the study are listed in **Table S2**, and details can be found in Supplementary Methods: *Construction of plasmids*.

### HPLC-based nucleotide quantification

*E. coli* strain BW2511324 and *E. coli* BW2511324 Δ*relA* was transformed with PanT_Bur. phage_ -expressing plasmid (pBAD33 - *Burkholderia* prophage phi52237) as well as empty pKK223-3 vector. The starter cultures were pre-grown overnight at 37 °C with vigorous shaking (200 rpm) in Neidhardt MOPS minimal (PMID:4604283) media supplemented with 1 µg/mL thiamine, 1% glucose, 0.1% caa, 100 µg/mL carbenicillin and 20 µg/mL chloramphenicol. The overnight cultures were diluted to OD_600_ 0.05 in 115 mL of pre-warmed medium MOPS supplemented with 0.5% glycerol as carbon source and grown until OD_600_ ≈ 0.2 at 37 °C, 200 rpm. At this point 0.2% arabinose was added to induce the expression of the toxin. 26 mL samples were collected for HPLC analyses at 0, 2, 5 and 10 minutes after the addition of arabinose and 1 mM IPTG. Nucleotide extraction and HPLC analyses were performed as described previously (23). The OD_600_ measurements were performed in parallel with collection of the samples for HPLC analyses.

### Toxicity neutralisation assays

Toxicity-neutralisation assays were performed on LB medium (Lennox) plates (VWR). *E. coli* BW25113 strains transformed with pBAD33 derivative plasmids encoding toxins (medium copy number, p15A origin of replication, Cml^R^, toxins are expressed under the control of a P_BAD_ promoter (36)) and pKK223-3 derivatives encoding antitoxins (medium copy number, ColE1 origin of replication, Amp^R^, antitoxins are expressed under the control of a P_Tac_ promoter (37)) were grown in liquid LB medium (BD) supplemented with 100 µg/mL carbenicillin (AppliChem) and 20 µg/mL chloramphenicol (AppliChem) as well as 1% glucose (repression conditions). Serial ten-fold dilutions were spotted (5 µL per spot) on solid LB plates containing carbenicillin and chloramphenicol in addition to either 1% glucose (repressive conditions), or 0.2% arabinose combined with 1 mM IPTG (induction conditions). Plates were scored after an overnight incubation at 37 °C.

To quantify bacterial viability (<u>C</u>olony <u>F</u>orming <u>U</u>nits, CFU), overnight cultures were diluted to OD_600_ either in the range from 0.1 to 0.01 (for the strains expressing PhRel2_*Bac. sub*._, with and without co-expression of wild-type PanA_*Vib. har*._) or OD_600_ ranging from 1.0 × 10^−4^ to 1.0 × 10^−5^ (all other strains) and spread on the LB agar medium as described above for the spot-test toxicity neutralisation assay. The final CFU/mL estimates were normalized to OD_600_ of 1.0.

### PanAT complex formation

Plasmids were transformed into *E. coli* BL21 DE3 strain. Fresh transformants were washed from an LB (Lysogeny broth, BD Difco–Fisher Scientific) agar plate and used to inoculate a 1 L culture LB supplemented with kanamycin (50 µg/mL). Cells were grown on 37 °C until OD_600_ reached 0.4-0.5 and induced with 0.5 mM IPTG. Cells were harvested after overnight cultivation on 18 °C, 220 rpm. Cells were opened with sonication in Binding Buffer (BB: 25 mM HEPES, pH 7.5; 300 mM NaCl; 10 mM imidazole; 2 mM CaCl_2_; 2 mM β-ME). Filtered lysate was incubated with 1 mL of previously buffer equilibrated Ni-beads (His60 Ni Superflow Resin, TaKaRa, Japan) for 30 minutes. Bound protein was washed with BB on gravity column and eluted with 300 mM imidazole. Fractions were resolved on 15% SDS-PAGE.

### Fluorescence microscopy

Overnight cultures were grown at 37 °C in MOPS minimal medium supplemented with 1% glucose. Next morning, the cells were washed, resuspended, and diluted in MOPS medium supplemented with 0.5% glycerol in order to remove glucose, followed by incubation at 37 °C until an OD_600_ of 0.3. For maintaining the plasmids, all cultures were grown in the presence of 34 µg/mL chloramphenicol. Toxin production was induced by addition of arabinose (0.2%) for 30 min, followed by staining with 200 nM of the membrane permeability indicator SYTOX Green (26) alongside the induction, and 250 nM of the membrane potential-sensitive dye DiSC_3_(5) (38) for the last 5 min (25, 39). Samples were immobilised on microscope slides covered with a thin layer of H_2_O/1.2% agarose and imaged immediately. As a positive control for pore-formation, BW25113 *E. coli* cells transformed with the empty pBAD33 vector were incubated with 10 µg/mL polymyxin B for 15 min (40). Microscopy was performed using a Nikon Eclipse Ti equipped with Nikon Plan Apo 100×/1.40 Oil Ph3 objective, CoolLED pE-4000 light source, Photometrics BSI sCMOS camera, and Chroma 49002 (EX470/40, DM495lpxr, EM525/50) and Semrock Cy5-4040C (EX 628/40, DM660lp, EM 692/40) filter sets. Images were acquired with Metamorph 7.7 (MolecularDevices) and analysed with Fiji (41).

### Selection of cross-inhibiting PanA mutants

An error-prone PCR mutant library of *Vibrio Harveyii* PanA antitoxins was created as described in Supplementary Methods: *Selection of cross-neutralising PanAs: preparation of the antitoxin mutant library*. Five µL (around one µg) of the antitoxin mutant library was transformed into the BW25113 *E. coli* strain carrying a non-cognate toxin expression plasmid PhRel2_*Bac. sub*._ toxSAS toxin from *B. subtilis* la1a (VHp303). The transformants were let to recover for one hour in 1 mL of SOC at 37 °C and added to 20 mL of LB media supplemented with ampicillin (100 µg/mL), chloramphenicol (25 µg/mL), 0.2% L-arabinose and 1 mM IPTG. The bacteria were grown overnight at 37 °C while expressing both toxin and antitoxin. Next day the plasmid was extracted from 3 mL of the culture using Favorprep Plasmid Extraction Mini Kit (Favorgen Biotech Corp.). 500 ng of the plasmid mix was again transformed into BW25113 carrying a toxin expression plasmid and let to recover as before. 100 µL of the recovery culture was spread on LB agar plates containing corresponding antibiotics as well as 0.2 % Glucose (control of transformation efficiency), and the rest of the culture was collected by centrifugation and spread on an LB agar plate containing corresponding antibiotics as well 0.2% L-arabinose and 1 mM IPTG.

Overnight cultures were started from selected colonies for further testing of cross-inhibition. Plasmids were extracted with Favorprep Plasmid Extraction Mini Kit and cleaved with FastDigest SacI restriction enzyme (Thermo Scientific) to eliminate the toxin plasmids. To ensure the purity of the antitoxin mutant mix it was transformed into *E. coli* DH5α strain and plasmids were extracted from the offspring of a single colony. 500 ng of the mutated plasmid was transformed into cognate or non-cognate toxin expressing *E. coli* BW25113 strain. Again, 100 µL of recovery culture was spread onto LB supplemented with corresponding antibiotics as well as 0.2% glucose agar plates, and rest of the bacteria were collected and spread LB agar plates supplemented with corresponding antibiotics, 0.2% arabinose and 1 mM IPTG. Phusion High-Fidelity DNA Polymerase (Thermo Scientific) was used to amplify the *panA* mutant (pK223_fwd_CPEC and pK223_rev_CPEC primers) and toxin genes (pBAD_fwd and pBAD_rev primers) with colony PCR and sequenced using pK223_rev_CPEC or pBAD_fwd primer correspondingly. Plasmid mixes and bacterial colonies were tested for possible contamination at various steps using FIREPol DNA Polymerase (Solis BioDyne): antitoxins were tested with the combination of pK223_rev_CPEC and STEC_panA_ctrl2, VH_panA_ctrl1 or Bsup_panA_ctrl1 primers, and toxins with the combination of pBAD_fwd and STEC_TOX_ctrl1, VH_TOX_ctrl1, Bsup_TOX_ctrl1 (**Table S2**).

## Acknowledgements

We are thankful to Tsigereda Ghebretnsae Ghebrelul and Constantine Stavropoulos for their help with microbiological assays, Niilo Kaldalu and Villu Kasari for sharing *E. coli* strain lacking Lon and ClpPX, and Michael Laub for sharing SocA/B-encoding plasmids. We are also grateful to Mikael Lindberg and the Umeå University Protein Expression Platform (PEP). This research was supported by grants from the Swedish Research Council (Vetenskapsrådet) grants (2017-03783 to V.H. and 2019-01085 to G.C.A.), Ragnar Söderbergs Stiftelse (to V.H.), postdoctoral grant from the Umeå Centre for Microbial Research, UCMR (to H.T.), the European Union from the European Regional Development Fund through the Centre of Excellence in Molecular Cell Engineering (2014-2020.4.01.15-0013 to T.T. and V.H.); and the Estonian Research Council (PRG335 to T.T. and V.H.). V.H. group was also supported by the Swedish Research Council (2018-00956 to V.H.) within the RIBOTARGET consortium under the framework of JPIAMR. H.S was supported by UK Biotechnology and Biological Sciences Research Council grant (BB/S00257X/1) and J.A.B by a UK Medical Research Council grant (MR/N013840/1.)

