## Supplementary figures and images for "Panacea: a hyperpromiscuous antitoxin protein domain for the neutralisation of diverse toxin domains"

### Dataset S2

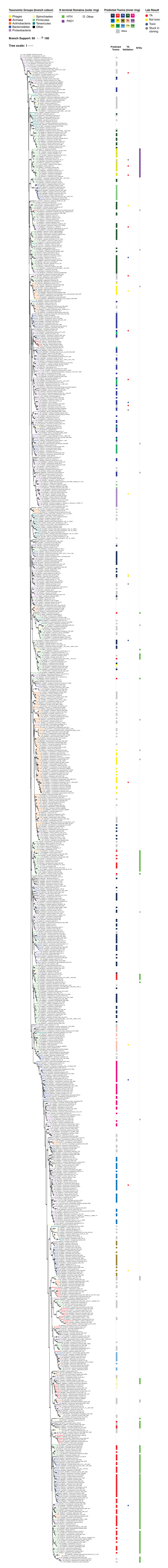
