## Supplementary material for "Panacea: a hyperpromiscuous antitoxin protein domain for the neutralisation of diverse toxin domains": SI Text and Figures

[†] equal contribution

[*] corresponding authors:

**Supplementary methods**

*Representative sequence dataset assembly*

To reduce the initial dataset of 2,281 DUF4065-containing sequences (sequence_set **1**) to a representative set for phylogenetic and gene neighbourhood figures, we used Usearch v11.0.667 (options: -cluster-fast, -id 0.5 or 0.1, -centroids) (1) to select sequences that are representatives of 50% identical and 10% identical clusters, respectively. The 50% resulted in 989 sequences (sequence_set **2**) and the 10% threshold resulted in 483 sequences (sequence_set **3**). All sequence sets are available online, along with their associated alignments (before and after trimming; see below) and where applicable, trees in Newick format (https://github.com/GCA-VH-lab/Panacea).

*Phylogenetic analysis*

Sequences were aligned with MAFFT as above, and alignment positions with >50% gaps were removed with trimAL v. 1.4.rev22 (2). Maximum Likelihood phylogenetic analysis was carried out with and IQ-TREEv1.6.10 (3) on the Cipres Science Gateway v 3.3 portal, with the best-fit model being determined by the program (4). The ultrafast bootstrapping (UFB) approximation method was used to compute support values for branches out of 1000 replicates. Annotated phylogenetic tree figures of sequence_set **2** were created with iTOL (5).

*Prediction of sequence features and structure*

We used NCBI's Conserved Domain Database batch search (6) and – for representatives - HHPred (7) with default settings to annotate domains other than DUF4065 that are present in this dataset. We made an alignment from sequence_set **2** using MAFFT v. 7.453 with the L-INS-i strategy (8), which allowed us to find a new N-terminal domain, which we named as PanA associated domain 1 (PAD1). The PAD1 domain has no identifiable homology with any domain in the NCBI CDD, or – with any probability greater than 50% – to any known structure that can be searched with HHPred (7). Using the NCBI CDD server, we identified 107 proteins from sequence_set **2** that have the HTH domain in their N-terminal region. We used MAFFT as above to align these 107 proteins and made an updated HTH HMM that was further used to identify HTH domain hits in sequence_set **1**. N-terminal extensions preceding the DUF4065 domain containing at least 80 amino acids which were not identified as HTH or PAD1 were named as 'Other' (**Table S1**) (9). Sequence conservation logos were made with WebLogo (10).

To detect if a PanA-encoding region is bacteriophage in origin, we used the tool PHASTER (11), to analyse a DNA segment equivalent to four upstream and four downstream genes around PanA from the 25 PanA-carrying taxa that have been tested in toxicity neutralisation assays.

*Prediction of TA loci*

Our Python tool FlaGs (12), which takes advantage of the sensitive sequence search method Jackhmmer (13), was adapted to identify conserved two- or three-gene conserved architectures that are typical of TA loci using the 2281 sequences (sequence_set **1**) as input. This Python script (available from https://github.com/GCA-VH-lab/Panacea) identifies two or three-gene architectures with up to 100 nucleotides between genes that are conserved in at least two different genomes, while the flanking regions are not conserved (in order to be sure of this, DUF4065-encoding genes were disregarded that were encoded at the end of contigs even if they otherwise appeared TA-like; **Dataset S1).** See **Figure S1** for details of the pipeline**.** Toxin clusters identified in this full dataset were recorded in **Dataset S1** and used to annotated the phylogenetic tree of sequence_set **2**.

To identify the most likely cases of false positives, where a panA gene happens to be proximal to non-toxin genes belonging to the same homologous cluster in two or more genomes purely by chance, we applied a reciprocality check, where each potential toxin hit was searched with BlastP (v. 2.9.0+; default settings) against the full proteome set, and the number of times the top five hits were found adjacent to PanA were scored (**Figure S1**, **Table S1**) . Finally, we identified third gene-encoded "accessory proteins" as those conserved in as a third gene in a subset of genomes that encode a particular predicted TA pair (**Fig. S1**, **Dataset S1**).

*Metabolic labelling*

Metabolic labelling assays were performed as described previously (14). One colony from freshly transformed (with a toxin or empty vector control) *E. coli* BW25113 cells was used in making overnight cultures, which were grown in defined Neidhardt MOPS minimal media (15) supplemented with 1% glucose, 0.1% of casamino acids and appropriate antibiotics at 37 °C with shaking overnight. The following morning, overnight cultures were diluted to an $OD_{600}$ of 0.05 in 15 mL MOPS minimal media with 0.5% glycerol as the carbon source, appropriate antibiotics, as well as a set of 19 amino acids (lacking methionine), each at final concentration of 19 µg/mL. The culture was grown until an $OD_{600}$ of 0.2-0.3 in a water bath with shaking (200 RPM) at 37 °C. The expression of toxins was induced with 0.2% L-arabinose. For a zero point, 1 mL of culture was taken and mixed with 2 µCi $^{35}$S methionine (Perkin Elmer), 0.65 µCi $^{3}$H uridine (Perkin Elmer) or 2 µCi $^{3}$H thymidine (Perkin Elmer) just prior to induction, while simultaneously another 1 mL of culture was taken for an $OD_{600}$ measurement. This was repeated with time points of 2, 5, 10 and 15 minutes. Incorporation of radioisotopes was halted after 8 minutes with addition of 200 uL of ice cold 50% trichloroacetic acid (TCA).

The resultant 1.2 mL culture/TCA samples were loaded onto GF/C filters (Whatman) prewashed with 5% TCA and unincorporated label was removed by washing the filter twice with 5 mL of ice-cold TCA followed by a wash with 5 mL 95% EtOH (twice). The filters were placed in scintillation vials, dried for at least 2 hours at room temperature, followed by the addition of EcoLite Liquid (MP Biomedicals) scintillation cocktail (5 mL per vial). After shaking for 15 minutes, radioactivity was quantified using TRI-CARB 4910TR 100 V scintillation counter (PerkinElmer).

*Construction of plasmids*

All bacterial strains and plasmids used in the study are listed in **Table S2**.

To test toxicity, toxin ORFs were cloned into the medium copy, arabinose inducible pBAD33 vector (16) between SacI and HindIII restriction sites. To make the constructs with a strong Shine-Dalgarno motif, the 5'-AGGAGG-3' sequence was incorporated into the pBAD33 vector. The full start codon context including the Shine-Dalgarno motif and intervening sequence was therefore 5'-AGGAGGAATTAAATG-3'. To test neutralisation of toxin, *panA* ORFs were cloned into the medium copy, IPTG inducible pKK223-3 vector between EcoRI and HindIII restriction sites. To test complex formation, cognate toxin and *panA* pairs were cloned as bicistronic operon into single pET24d vector (high copy, T7 promoter-driven expression) (Novagen). To express both toxin and PanA protein, toxin ORFs were cloned on native reading flame in pET24d vector with His tag, and tag-less *panA* was cloned downstream of toxin with the spacer 5'-TAAGCTTATAAGGAGGAAAAAAAA-3' includes the strong Shine-Dalgarno motif.

The plasmids were constructed with ligation, Circular Polymerase Extension Cloning (CPEC), Gibson assembly or site-directed mutagenesis method, materials and cloning methods are summarized in **Table S2**. Except for VHp680, VHp694, VHp709, VHp712 and VHp713, the plasmids used for toxicity neutralisation assays were constructed by the Protein Expertise Platform Umeå University (PEP). The toxin and *panA* ORFs were synthesised by Eurofins, amplified using primers containing the restriction sites consistent with cloning site in the vector and ligated with the vector linearised by the same set of restriction enzymes with each ORF. For VHp585 (pBAD33-SD-*socB*), pET22b-*socB* gifted by Michael Laub was used for amplification of *socB* ORF. For the other plasmids constructed by ligation, the ORFs were cloned by same way as with PEP, using primers, templates and vectors shown in **Table S2.** To construct plasmids by Gibson assembly, two or three DNA fragments per plasmid were amplified with the primers and templates shown in **Table S2** and assembled by NEBuilder HiFi DNA Assembly Cloning Kit (NEB). For site-directed mutagenesis to construct the plasmids coding individual *panA*$_{\textit{Vib. har.}}$ mutants, the whole VHp474 plasmid was amplified using 5' phosphorylated-primers (**Table S2**) containing the mutation. After DpnI treatment, the PCR fragment was self-circulated by blunt-end ligation. Ligation or assembly mixes were transformed by heat-shock in *E. coli* DH5$\alpha$. PCR amplifications were carried out using Phusion polymerase, purchased from ThermoScientific along with restriction enzymes, T4 PNK and T4 ligase. Plasmid construction using CPEC is detailed in the following section.

*Selection of cross-neutralising PanAs: preparation of the antitoxin mutant library*

An error prone PCR library was generated using GeneMorph II Random Mutagenesis Kit (Agilent). VH_fwd_CPEC and VH_rev_CPEC primers (**Table S2**) were used to amplify the *panA* $_{\textit{Vib. har.}}$ gene of *Vibrio harveyi* from 1 ng of VHp474 plasmid with 32-cycle PCR run. The DNA fragments were purified from agarose gel after electrophoresis using Zymoclean Gel DNA Recovery Kit (Zymo Research). Circular Polymerase Extension Cloning (CPEC (17)) was used to insert the mutated fragments into pKK 223-3 plasmid. The vector was linearized using FastDigest EcoRI and HindIII restriction enzymes (Thermo Scientific), run on an agarose gel and purified with Zymoclean Gel DNA Recovery Kit. 500 ng of linearized vector and 120 ng of PCR fragment (1:2 molar ratio) was used for one cloning reaction (50 µL, 25 cycles, Phusion High-Fidelity DNA Polymerase by Thermo Scientific). 15 CPEC reactions were pooled together and purified using DNA Clean & Concentrator Kit (Zymo Research). 2.5-5 µL of the plasmid library was transformed into either DH5α chemical competent cells prepared using Mix & Go! *E. coli* Transformation Kit (Zymo Research) or into DH5α electroporation competent cells prepared in-house. 1 mL of prewarmed SOC medium was added to transformants and they were let to recover for one hour on 37 °C with shaking. Recovery culture was added to 11 mL of prewarmed LB supplemented with ampicillin (100 µg /mL final concentration) and grown overnight at 37 °C. Plasmids were extracted from 3 mL of overnight culture using Favorprep Plasmid Extraction Mini Kit (Favorgen Biotech Corp.) and pooled together. The size of the library was assessed from plating serial dilutions of recovery culture on LB-agarose plates supplemented with ampicillin (100 µg /mL final concentration; estimated library size 300,000 mutants).

**Table S1. Summary of experimental validation and *in silico* prediction of novel PanAT pairs that were not experimentally investigated in detail.**

| | **Experimental validation** | | | | |
|---|---|---|---|---|---|
| **Organism** | **toxin** | **antitoxin** | **Predicted toxin MOA / toxin family** | **Toxin accession** | **Antitoxin accession** |
| *Planomicrobium flavidum* | **stuck in DNA cloning** | n/a | toxSAS | WP_088005586.1 | WP_088005585.1 |
| *Pararheinheimera mesophila* | **stuck in DNA cloning** | n/a | membrane | WP_046521261.1 | WP_046521262.1 |
| *Neisseria gonorrhoeae* | **stuck in DNA cloning** | n/a | mRNAse | WP_047918230.1 | WP_003695998.1 |
| *Clostridium cadaveris* | **toxic** | failed to neutralise | mRNAse | WP_074844970.1 | WP_074844973.1 |
| *Streptococcus* phage P9 | **toxic** | failed to neutralise | membrane | YP_001469232.1 | YP_001469231.1 |
| *Vibrio cholerae* | **toxic** | toxic | membrane | WP_088132352.1 | WP_088132353.1 |
| *Herbaspirillum frisingense* | **toxic** | toxic | MqsR | WP_006465653.1 | WP_006465652.1 |
| *Chryseobacterium bovis* | **toxic** | toxic | MazF | WP_076782181.1 | WP_076782182.1 |
| *Bifidobacterium adolescentis* | not toxic | n/a | Fic/Doc | WP_003806793.1 | WP_003806794.1 |
| *Clostridioides difficile* | not toxic | n/a | membrane | WP_074138355.1 | WP_074138354.1 |
| *Escherichia coli* MOD1 | not toxic | n/a | mRNAse | WP_077635758.1 | WP_089622957.1 |
| *Campylobacter jejuni* | not toxic | n/a | VopL-like | WP_002900950.1 | WP_002900952.1 |
| *Salmonella* phage vB SemP Emek | not toxic | n/a | mRNAse | YP_006560566.1 | YP_006560565.1 |
| *Flavobacterium xanthum* | not toxic | n/a | MazF | WP_073354223.1 | WP_084129631.1 |
| *Listeria* phage B025 | not toxic | n/a | mRNAse | YP_001468667.1 | YP_001468666.1 |
| *Caulobacter crescentus* | not toxic | n/a | SocB | WP_010921343.1 | WP_010921344.1 |

### Supplementary Figures

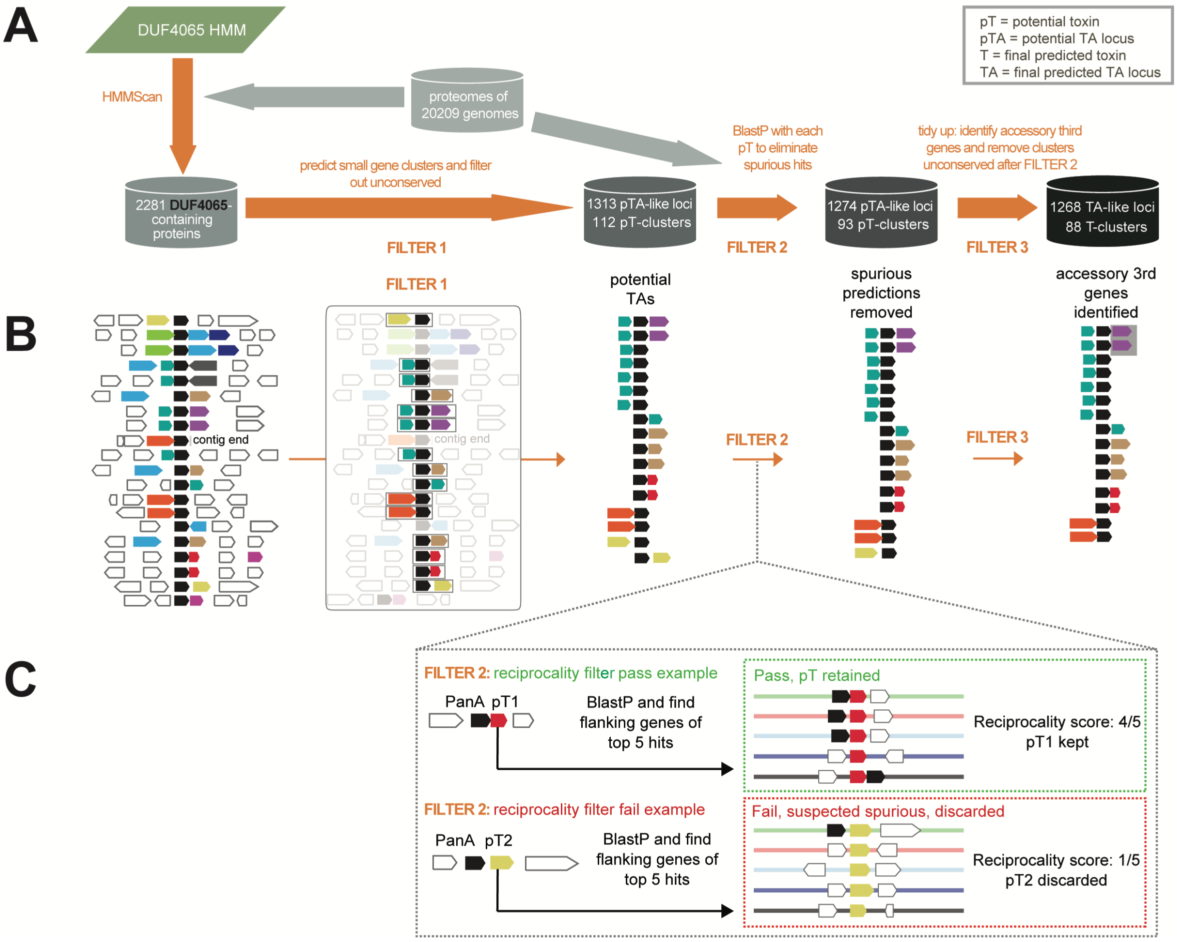


**Figure S1. The bioinformatic workflow for TA prediction.**

(**A**) The overall workflow. To detect the presence of PanA across the tree of life we used the Hidden Markov Model (HMM) of the DUF4065 domain from Pfam database which is carried by PanA. We scanned our database of proteomes with the DUF4065 HMM using thresholds set to the HMM profile's gathering cutoffs. We found that the DUF4065 domain was present in 2,281 identified sequences. After identifying conserved clusters of flanking genes and filtering, we predicted 1,268 individual TA-like operons, with toxins that can be clustered into 88 conserved clusters.

(**B**) Two- or three-gene conserved architectures that are typical of TA loci were predicted using the 2281 sequences as input. In total, we predicted 1,313 preliminarily TA (pTA)-like loci, using the filtering criteria:

i) that there should be a maximum distance of 100 nucleotides between the two genes,

ii) that this architecture is conserved in two or more species and

iii) the conservation of the gene neighbourhood does not suggest longer operons than three genes. In order to be sure of this, DUF4065-encoding genes were disregarded that were encoded at the end of contigs even if they otherwise appeared TA-like.

The second filter was designed to remove the most likely cases of spurious hits (**C**, see below). In the third filter we identified third gene-encoded "accessory proteins" as those conserved in as a third gene in a subset of genomes that encode a particular predicted TA pair.

(**C**) Further detail of the second filter. To identify the most likely cases of false positives, where a *panA* gene happens to be proximal to non-toxin genes belonging to the same homologous cluster in two or more genomes purely by chance, we applied a reciprocality check, where each potential toxin hit was searched with BlastP against the full proteome set, and the number of times the top five hits were found adjacent to PanA were counted. This gave a score out of five for the association of this putative toxin with PanA. For example, a score 1/5 means that that putative toxin is only adjacent to PanA once out of its top five relatives, and is therefore not reliable as a TA prediction. The best possible score 5/5 on the other hand means that all the five top hits are in the same TA-like locus with PanA, and therefore the association of the putative toxin and PanA has full reciprocity. Only pairs with a score greater than 1 pass this filter.

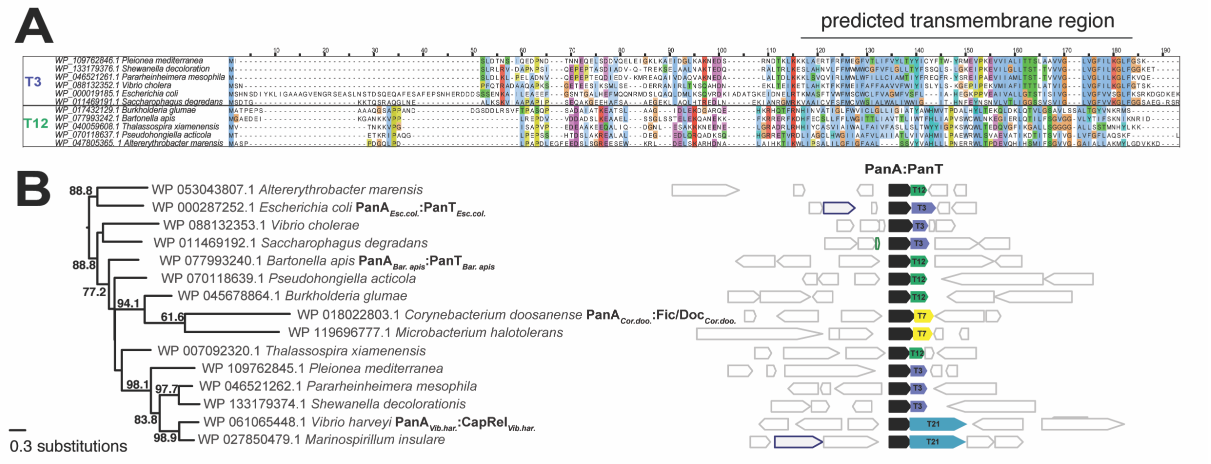


**Figure S2. Toxin partner switching in a branch of the PanA tree, panel B is related to Figure 1.**
(**A**) Multiple sequence alignment shows the two verified toxin clusters T3 and T12 with transmembrane domains are distinct, but are potentially homologous.
(**B**) The identity of the PanT toxin does not follow the phylogeny of the PanA partner. While T12 and T3 are potentially structurally related membrane-interacting proteins, T7 (Fic/Doc) and T21 (ToxSAS) are structurally unrelated.

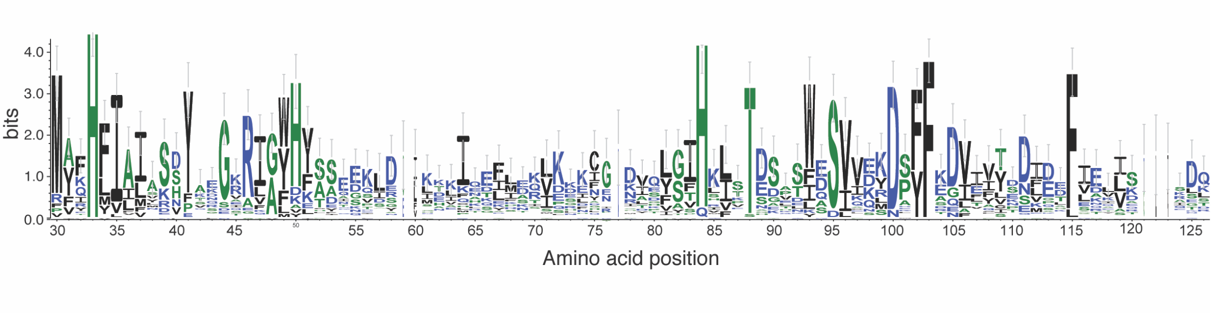


**Figure S3. Sequence logo of the PAD1 domain shows conserved amino acids.**
The logo was made from an alignment of 33 detected PAD1 sequence regions using the WebLogo server (10).

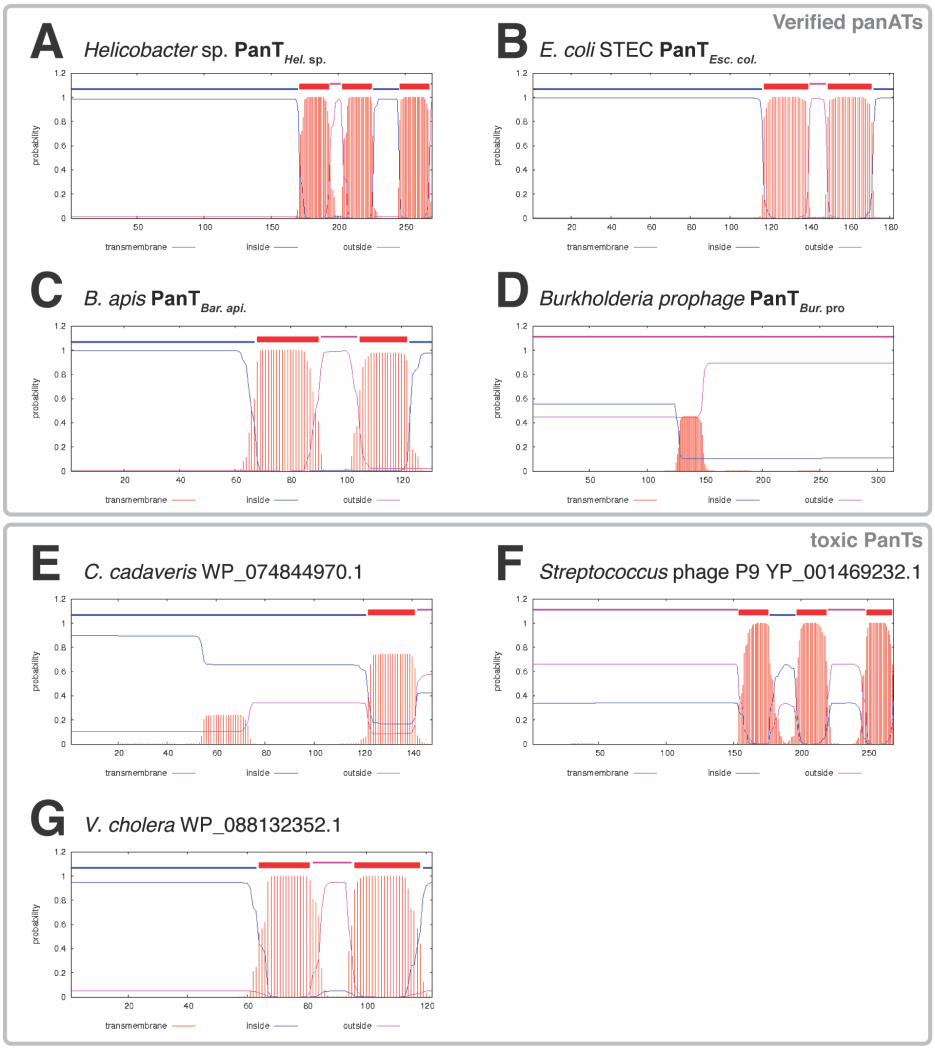


**Figure S4. Seven identified PanT toxins carry transmembrane regions, related to Figure 5 and Tables 1 and S1.**
Transmembrane predictions of four toxins from verified PanATs (**A-D**), and three toxins where the PanA did not neutralise the cognate PanT in our experimental system (**E-G**). Transmembrane regions were predicted with the TMHMM 2.0 sever (9).

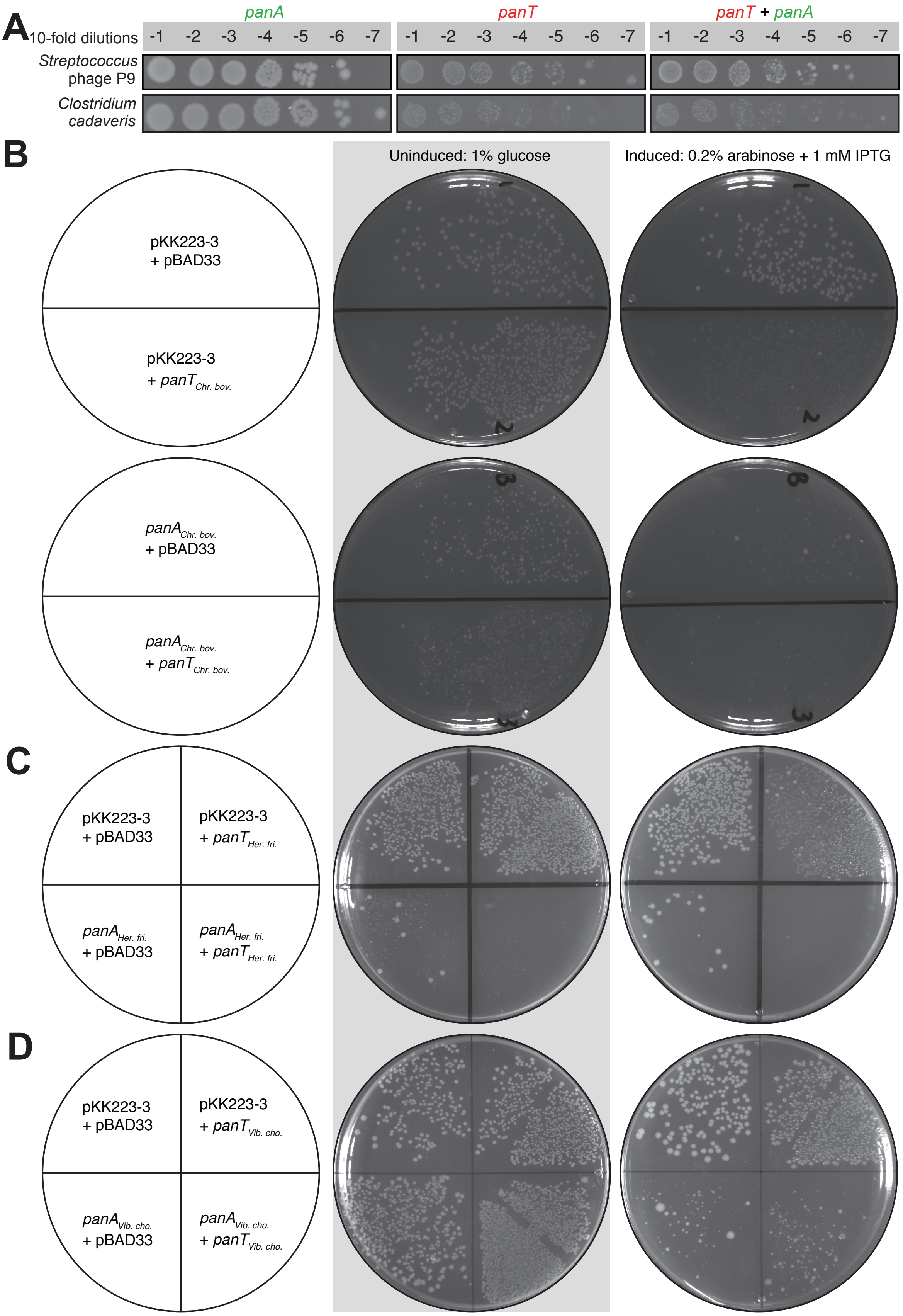


**Figure S5. When expressed *E. coli* BW25113S some of the PanAT representatives do not act as typical TA pairs, related to Figure 1 and Table S1.**

(**A**) *Streptococcus* phage P9 and *Clostridium cadaveris* PanA representatives fail to counteract the toxicity of cognate PanTs. Overnight cultures of *E. coli* strains transformed with pBAD33 and pKK223-3 vectors or derivatives expressing putative *panT* toxins and *panA* antitoxins, correspondingly, were adjusted to $OD_{600}$ 0.1, serially diluted from $10^1$- to $10^7$-fold and spotted on LB medium supplemented with appropriate antibiotics and inducers (0.2% arabinose for *panA* induction and 1 mM IPTG for *panT* induction).

(**B**-**D**) Overexpression of (**B**) *Chryseobacterium bovis*, (**C**) *Herbaspirillum frisingense* and (**D**) *Vibrio cholerae* PanA representatives is toxic to *E. coli*. *E. coli* BW25113 was co-transformed with both pBAD33-based plasmid (either empty vector or L-arabinose inducible *panT* expression plasmid) and pKK223-3-based plasmid (either empty vector or IPTG-inducible *panA* expression plasmid), the cells were directly spread on uninducing (1% glucose) or inducing (0.2% arabinose and 1 mM IPTG) LB plates supplemented with appropriate antibiotics. The plates were scored after an overnight incubation at 37 °C. In the case of *C. bovis* and *H. frisingense* PanA representatives, even leaky expression of *panA* is toxic.

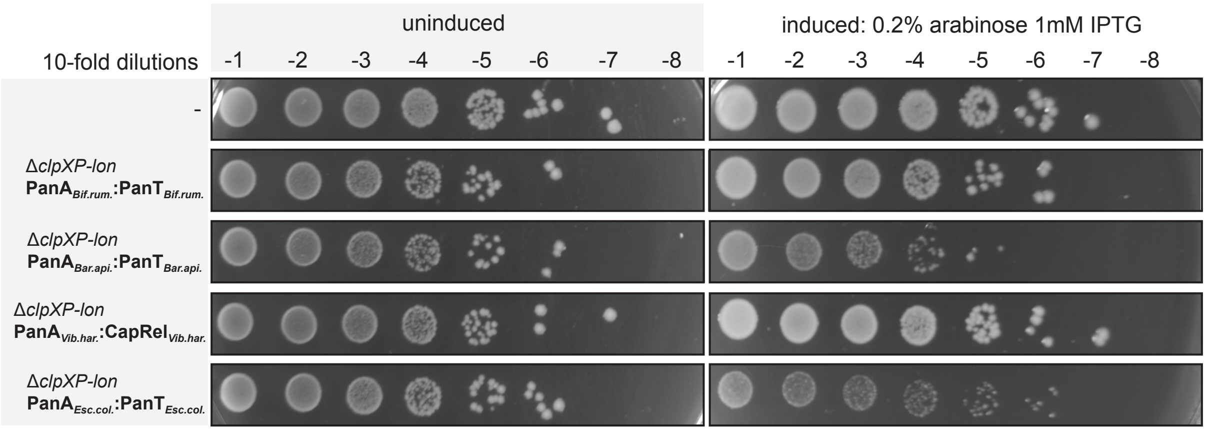


**Figure S6. PanA antitoxins can efficiently counteract the toxicity of cognate PanT in protease-deficient *E. coli*.**
Overnight cultures of *E. coli* strains transformed with pBAD33 and pKK223-3 vectors or derivatives expressing putative *panT* toxins and *panA* antitoxins, correspondingly, were adjusted to $OD_{600}$ 0.1, serially diluted from $10^1$ to $10^7$-fold and spotted on LB medium supplemented with appropriate antibiotics and inducers (0.2% arabinose for *panA* induction and 1 mM IPTG for *panT* induction).

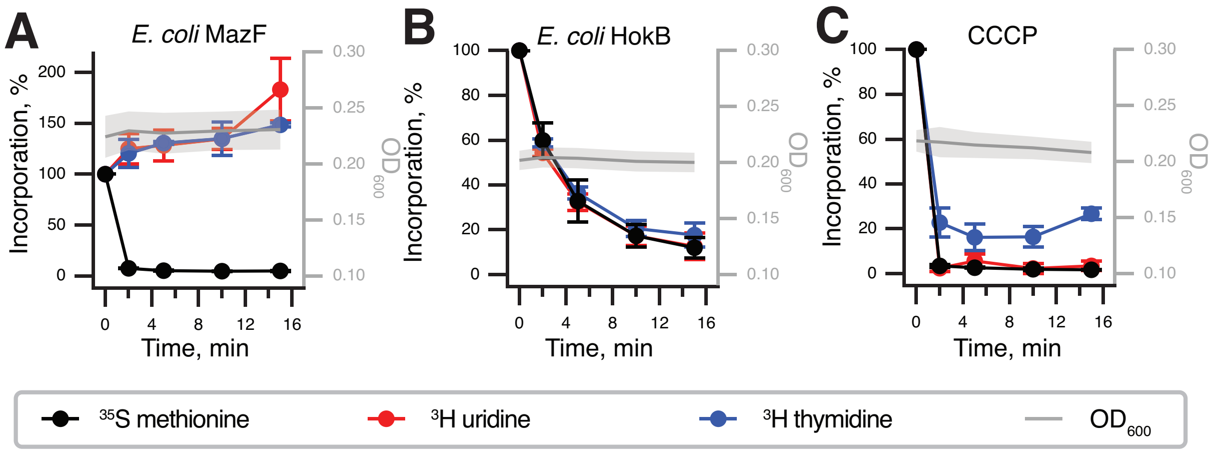


**Figure S7. Metabolic labelling assays with well-characterised known *E. coli* TA toxins and CCCP membrane disruptor, related to Figures 3, 4 and 5.**
Metabolic labelling assays were performed with wild-type *E. coli* BW25113 expressing (**A**) *mazF,* (**B**) *hokB* or wild-type *E. coli* BW25113 treated with CCCP (**C**).

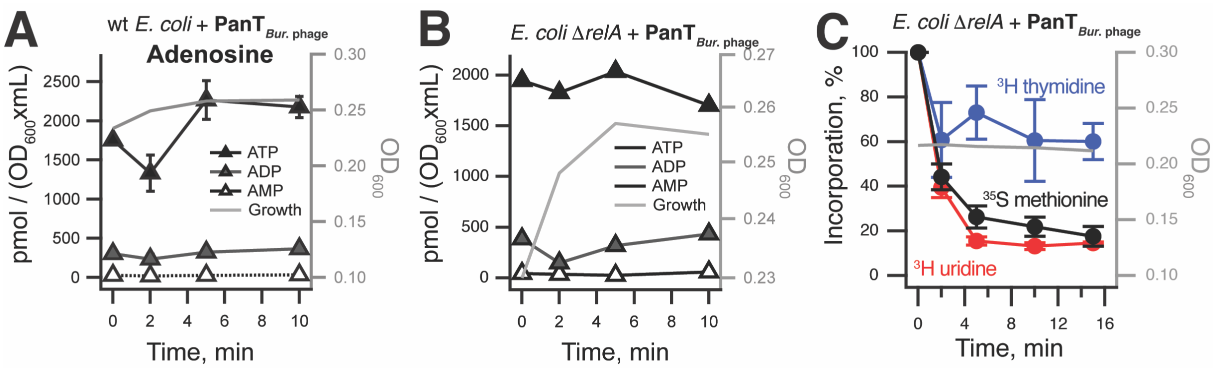


**Figure S8. PanT$_{Bur.\ phage}$ toxin does not induce the stringent response in absence of RelA, related to Figure 4.**
(**A**, **B**) Nucleotide pools of wild-type (**A**) or Δ*relA* (**B**) *E. coli* BW25113 expressing PanT$_{Bur.\ phage}$ toxin. Cell cultures were grown in defined minimal MOPS medium supplemented with 0.5% glycerol at 37 °C with vigorous aeration. The expression of PanT$_{Bur.\ phage}$ toxin was induced with 0.2% L-arabinose at $OD_{600}$ 0.2. Intracellular nucleotides are expressed in pmol per $OD_{600}$ • mL as per the insert. Error bars indicate the standard error of the arithmetic mean of three biological replicates. (**C**) Metabolic labelling assay using Δ*relA E. coli* BW25113 expressing PanT$_{Bur.\ phage}$ toxin.

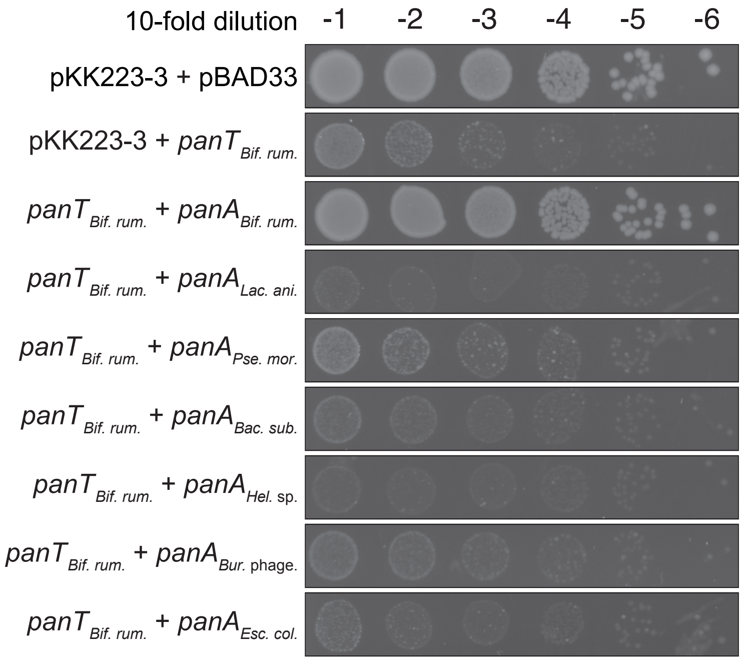


**Figure S9. Neutralisation of wild-type *panT*$_{\textit{Bif. rum.}}$ by a selection of *panA* antitoxins, related to Figure 6A.**
The overnight cultures of *E. coli* strains transformed with pBAD33 and pKK223-3 vectors or derivatives thereof expressing toxin and PanA antitoxins, was adjusted to 1.0, cultures serially diluted from $10^1$- to $10^7$-fold and spotted on LB agar medium supplemented with appropriate antibiotics as well as inducers (0.2% arabinose for toxin induction and 1 mM IPTG for induction PanA variants). The plates were scored after for 14 hours at 37 °C. The $10^{-1}$ dilution spots are reproduced on **Figure 6A**.

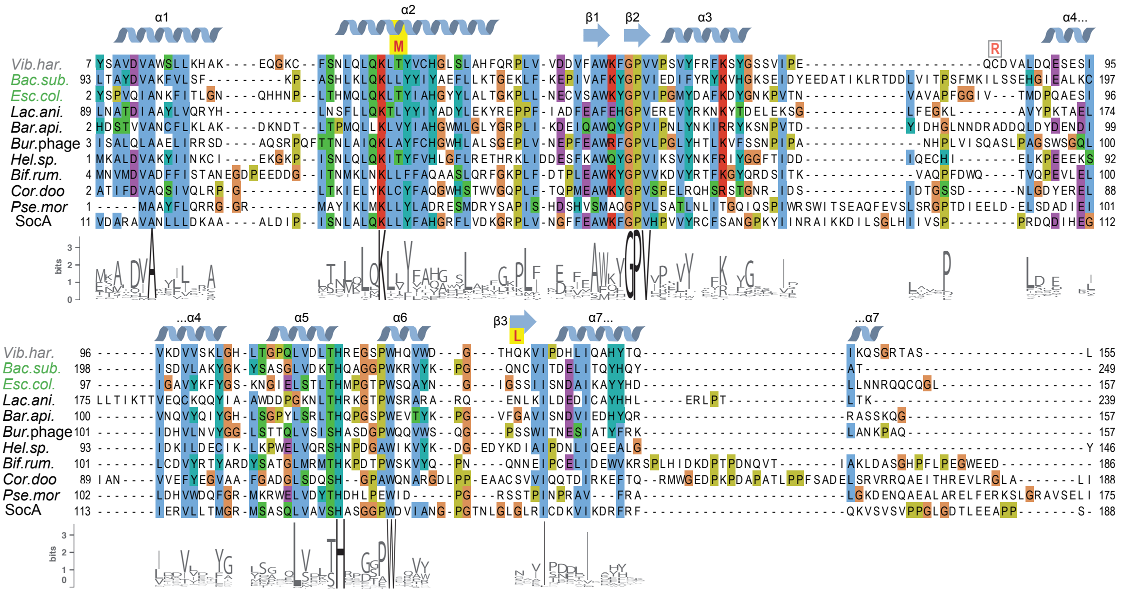


**Fig S10. Sequence alignment of Panacea domains of verified PanAs and SocA, related to Figure 6.**

N-terminal regions, including the PAD1 domains of toxSAS toxins are not shown. The predicted secondary structure of PanA*Vib. har.* is shown above the alignment. Sequence conservation logos computed from the **Figure 1** PanA dataset are shown below the alignment. Amino acid substitutions in PanA*Vib. har.* that allow cross-neutralisation are shown above the alignment in red, highlighted with yellow boxes.

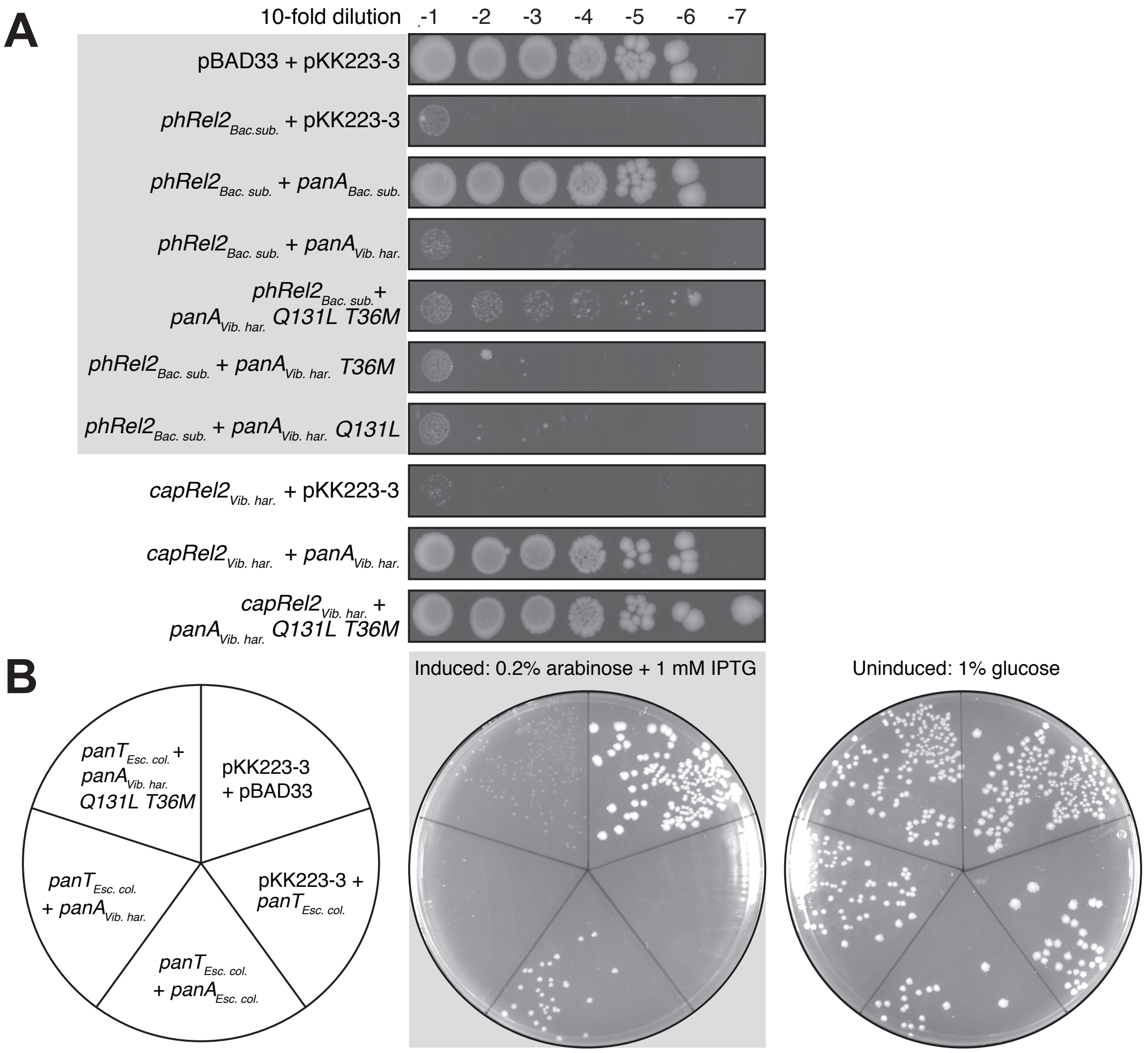


**Figure S11. PanA*Vib. har.* T36M Q131L neutralises both PhRel2*Bac. sub.* and PanT*Esc. col.* but the protection is less efficient than in the case of cognate PanA*Bac. sub* and PanA *Esc. col.* antitoxins, related to Figure 6.**

(**A**) Toxicity neutralisation PhRel2*Bac. sub.* and CapRel*Vib. har.* by PanA*Vib. har.* T36M Q131L variant. Overnight cultures of *E. coli* strains transformed with pBAD33 and pKK223-3 vectors or derivatives expressing putative *panT* toxins and *panA* antitoxins, correspondingly, were adjusted to $OD_{600}$ 0.1, serially diluted from $10^1$ to $10^7$-fold and spotted on LB medium supplemented with appropriate antibiotics and inducers (0.2% arabinose for *panA* induction and 1 mM IPTG for *panT* induction). The plates were scored after 48 hours incubation at 37 °C.

(**B**) Toxicity neutralisation PanT*Esc. col.* by PanA*Vib. har.* T36M Q131L variant. After transformation, the cells were directly spread on uninducing (1% glucose) or inducing (0.2% arabinose and 1 mM IPTG) LB plates supplemented with appropriate antibiotics; scored after for 48 hours at 37 °C.
